# Goal-dependent resource-rational compression of attribute differences explains nonlinearities in multi-attribute decision making

**DOI:** 10.64898/2026.06.10.731311

**Authors:** Sherry Dongqi Bao, Saurabh Bedi, Da Li, Christian C. Ruff, Todd A. Hare

**Author notes:** These authors contributed equally to this work (Sherry Dongqi Bao, Saurabh Bedi).

## Abstract

Why do multi-attribute choices so often depart from classical weighted-additive decision rules? Rather than attributing such deviations solely to biases or heuristics, we propose a resource-rational account in which value differences are encoded via capacity-limited information channels. Under resource-rational compression, these difference representations are systematically distorted, such that behavior deviates from weighted-additive predictions because value differences are not represented veridically. This theoretical account makes testable predictions about power-law relationships between true and internally represented differences. The amount of power-law-like compression is determined by information-processing capacity, emergent long-tailed prior distributions over attribute differences, and, in choice contexts, goal-dependent subjective weights that govern the allocation of limited capacity across attribute channels. We test and find support for these predictions in an attribute difference-estimation task and by reanalyzing existing food- and social-choice datasets. These results provide converging evidence for a normative, information-theoretic account of systematic nonlinearities in multi-attribute decision making. Together they show how goals can interact with cognitive capacity and priors to shape representational precision in ways that may facilitate or impair decision making.

## Introduction

In everyday life situations, people often face the challenge of choosing between options that differ in the advantages and disadvantages of their various attributes. For example, they may have to choose between foods that are healthier but less flavorful and foods that are tastier but less healthy, or between an action that yields high personal benefits but also does substantial harm to others and an action with more equitable consequences.

Theories of preference-based decision making have commonly assumed that people compute option utilities based on stable, accurate value representations that are not systematically affected by information processing-resource constraints [1]. A canonical instantiation is the weighted-additive decision rule (WAD) for multi-attribute choice: decision makers multiply each attribute’s veridical value by a subjective weight reflecting its importance and then integrate these weighted values to obtain an overall option value [2]. The WAD is often used as the default model for multi-attribute decisions and is widely applied in individual and social choice contexts [3–6]. However, it is also well known that human behaviour frequently deviates from the predictions of decision models based on canonical weighted-additive decision rules [7]. To account for these shortcomings, numerous alternatives to WAD have been proposed to describe multi-attribute decisions, often proposing that behaviour is subject to irrational biases or effort-reducing heuristic simplifications [8–11]. Here, we show that deviations from WAD can emerge from normative computations under neuro-cognitive resource constraints, without invoking biases or heuristics.

Specifically, we propose that some such deviations arise because attribute values are not encoded veridically, but through resource-rational compressed representations. Our proposal builds on well-established findings in perceptual neuroscience showing that the brain, as a limited-capacity system, forms resource-rational compressed representations tuned to environmental structure and task demands [12–18]. We extend this resource-rational compression to multi-attribute valuation and propose that value difference representations needed for binary choice are distorted as a consequence of capacity limitations and environmental structure (priors). Critically, our goal-adaptive difference encoding account does not require a specific assumption about the distribution of individual attribute values. What is required is only that the task-relevant comparison variable (the difference between two attribute values that have to be compared for choice) has a long-tailed prior: small differences are encountered more often than large differences. A long-tailed prior, often referred to as a decaying prior, over differences emerges robustly from many distributions over individual values, including increasing, decreasing, and approximately Gaussian distributions, as shown in Supplementary Figure S1. Under the conditions of finite capacity and long-tailed priors, resource rational encoding produces compressed, approximately power-law mappings from objective to subjective differences. In both estimation and choice tasks, the degree of compression is determined by channel capacity and prior dispersion, while subjective weights over attributes also influence the strength of compression in choice tasks. This can make behavior appear to violate classical WAD predictions even when the underlying computation is optimal under these representational constraints.

We provide evidence in line with the goal-adaptive difference encoding model’s predictions from an attribute difference–estimation task and by reanalyzing existing multi-attribute choice datasets in the domains of food and social decision-making. The estimation task directly tests the prediction that subjective reports of attribute-value differences should show nonlinear compression relative to objective differences computed from participants’ earlier ratings. The estimation task also allows us to compare compression levels across attribute types while exogenously controlling the value of precisely representing each attribute. Specifically, we paid participants the same amount for accurately reporting each attribute difference and difference estimations did not imply selection or consumption of either item (i.e., subjective preferences were irrelevant). The choice analyses then ask whether the same resource-rational compression signature appears when attribute value differences are used to guide subjective decisions rather than explicit reports. Specifically, when analyzing the choice data we augment standard choice models with flexible nonlinear transformations of attribute differences and compare them to the canonical linear versions. In line with the goal-adaptive difference encoding account’s predictions, model comparison reveals attribute-specific degrees of nonlinearity consistent with capacity-limited encoding of attribute differences under stable and emergent decaying priors over value differences. Moreover, in choice, subjective importance weights can redistribute resources across attribute channels, producing a different ordering of relative compression (degree of nonlinearity) across attributes than observed in the estimation task. Lastly, we conduct an initial test of the role of capacity limits in the internal representation of attribute differences by comparing the degree of compression estimated in choice conditions with versus without time constraints (i.e., lower or higher capacity limits).

These empirical findings support our proposed normative, resource-rational explanation for systematic deviations from WAD-based ideal-observer predictions in multi-attribute decision making. Overall, the current work underscores the importance of accounting for representation-level constraints in multi-attribute valuation and choice. A mechanistic understanding of these constraints and their consequences allows for the design of targeted manipulations that increase or reduce the constraints to further test the theory and to develop interventions and procedures to support decision making in daily life.

## Results

### Theoretical account: goal-dependent encoding under capacity constraints

Building on recent information-theoretic accounts of perceptual biases [12, 13, 19], we propose a model of optimal encoding of attribute differences under capacity constraints. This model is aimed at explaining empirically observed nonlinear distortions in representations of attribute value differences in both attribute-difference estimation and multi-attribute choice tasks. In so doing, it also provides a principled account of why limited-capacity agents may systematically deviate from the predictions of classical weighted-additive decision rules.

The two task contexts we examine here share many features but there is also a key difference between them. In an attribute difference estimation task context, participants report the difference between two options on one or more specific attribute dimensions. In such a context, the goal is to represent and report the requested attribute value difference as accurately as resources allow. *The other attributes an object may contain and the participant’s subjective preferences over the different attributes are irrelevant in the estimation task*. In contrast, in a multi-attribute choice task, attributes differ in their subjective importance, such that estimation errors along certain attribute dimensions are more likely to result in undesirable outcomes than errors along other dimensions. The *weight or importance given to each attribute depends on the decision-maker’s goals and preferences*, and as we will show later, *determines how limited capacity is allocated across attribute channels, leading to goal-dependent levels of encoding precision* for the difference between options on a given attribute dimension.

In what follows, we characterize how a resource-bounded agent optimizes task performance by balancing the cost of information processing with limited resources against the benefit of achieving task goals. We show that such an information-theoretic constrained optimization solution leads to compressed, power-law-like relationships between true attribute value differences and their internal representations in both estimation and multi-attribute choice tasks. Moreover, this nonlinear relationship changes depending on the severity of resource constraints and the stimulus prior in the context of an estimation task. In a choice task, it is further modulated by goal-dependent, subjective attribute weights.

Note that because the precise shape of the power-law function jointly depends on multiple factors, including the capacity limits and the prior distribution function), we do not directly fit the individual parameters of the goal-dependent encoding model to human choice or estimation task data. Rather, we test for the presence of the power-law-like relationships the model predicts and if those relationships are altered in expected ways by experimental manipulations of task goals or resource constraints.

### Predicted representations within an estimation task

We first consider how attribute differences should be represented during an estimation task in which participants are asked to report the difference between two options along a single attribute dimension — such as the difference in the tastiness or healthfulness dimension between two food options. Representational limits imply that the agent does not have direct access to the true value difference *d*_*i*_ for attribute *i*, but instead forms an approximate internal representation through encoding. We formulate the resource-rational encoding over within-attribute differences, because the comparison signal *d*_*i*_ is the task-relevant input to estimation and is necessary for determining the attribute’s contribution in choices. At the same time, we treat individual attribute values as upstream inputs that may or may not already be distorted. We return to alternative possibilities and their implications in the Discussion.

Specifically, we let *r*_*i*_ denote the internal representation of the attribute value difference, and assume that it is drawn from a conditional encoding distribution *r*_*i*_ ∼ *Q*(*r*_*i*_ | *d*_*i*_).

The agent’s goal in the estimation task is to report value differences as accurately as possible. This corresponds to a loss function that minimizes expected squared error between the internal representation and the true value difference:

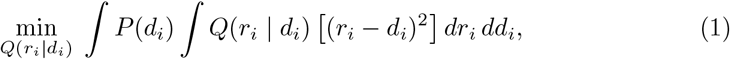

where *P* (*d*_*i*_) is the environmental distribution of the true value differences for attribute *i*, which is also a prior on how often the value difference *d*_*i*_ is encountered.

If the agent had infinite resources, the optimal representation would perfectly correspond to the true value difference, leading to noiseless representations and no behavioral variability or biases. However, neural information-processing capacity is constrained [20, 21], so that attribute channel *i* has finite capacity *C*_*i*_, and representations diverging too far from the prior for representations are costly [12, 13]. This constraint on information-processing capacity can then be formalized as a bound *C*_*i*_ on the Kullback–Leibler (KL) divergence [22] between the encoding distribution *Q*(*r*_*i*_ | *d*_*i*_) and the prior *P* (*r*_*i*_) [13, 19],:

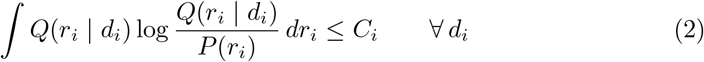

The solution to this constrained optimization yields the optimal encoder (see Methods):

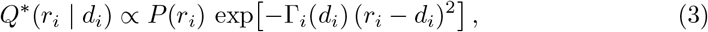

where the effective precision term 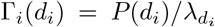 captures the joint influence of the environmental prior and the information-processing constraint. The Lagrange multiplier 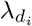 is determined by the capacity bound *C*_*i*_. It emerges from the constrained optimization, with larger *C*_*i*_ yielding smaller 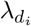 and less compression. For clarity of exposition, we refer to the effective capacity *C*_*i*_ as the key parameter governing compression.

We assume that the prior distribution of true value differences follows a power-law form,

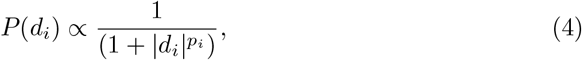

reflecting that small attribute differences are more frequent than large ones. We adopt this specific functional form for two reasons. First, across a wide range of priors that individuals might place on individual attribute values, the emergent distribution of pairwise differences is consistently long-tailed and decays for higher differences (see Fig. S1). Second, this form is a well-established assumption in models of numerosity estimation [13], which is conceptually fitting given that we represent attribute value differences as numerical magnitudes. (This form, with a typical exponent *p*_*i*_ = 2, is consistent with empirical regularities in number-word frequency [23, 24] and natural decision statistics [25], and is commonly used in resource-rational models of numerosity estimation [13].) Nevertheless, note that qualitatively similar predictions arise even when the prior over differences is shifted away from zero, as with truncated normal or truncated Cauchy distributions centered at nonzero values (see Fig. S2 and Fig. S3).

Variations in the power parameter *p*_*i*_ correspond to broader (smaller *p*_*i*_) or narrower (larger *p*_*i*_) priors, representing differences in environmental uncertainty that can modulate compression alongside capacity limits. Importantly, these compression patterns generalize across a wide range of monotonic, decaying priors, as shown by results for different degrees of prior dispersion (Figure 1).

**Fig. 1.**
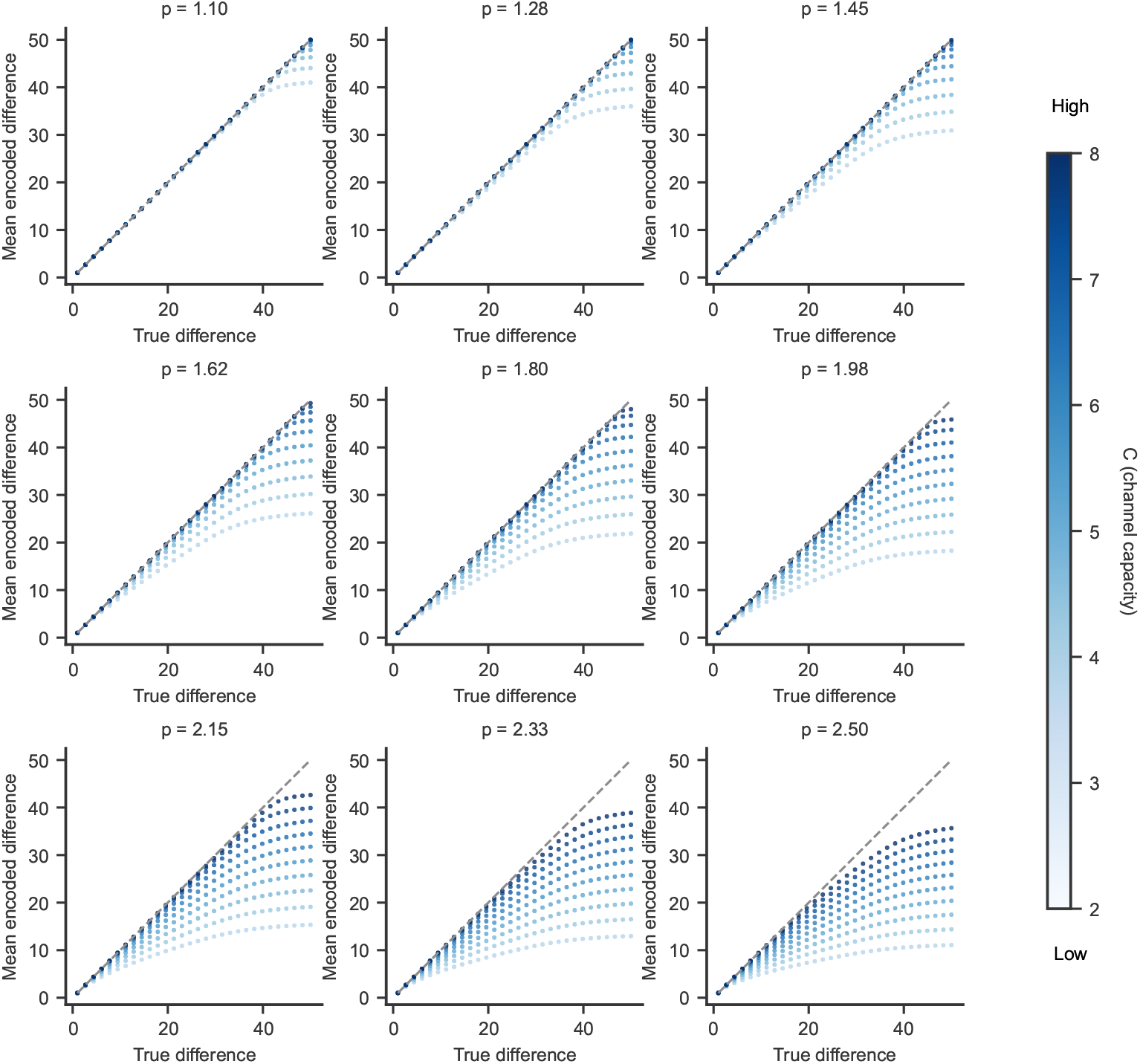
Channel capacity and prior dispersion jointly determine compression in an estimation task. Simulations for channel *i* follow the encoder formulated in Equation 3. The expectation 𝔼 [*r*_*i*_ | *d*_*i*_] provides the mean encoded difference responses shown along the *y*-axis. Within each panel, we vary the channel capacity parameter *C*, while keeping the prior fixed to a power-law form *P* (*d*) *∝* 1*/*(1 + |*d*|^*p*^). Higher *C* corresponds to higher capacity and produces responses closer to the identity line (exponent closer to 1 in the fitted power function). We used 10 evenly spaced values of *C ∈* [2, 8] for the simulation. Larger *p* yields narrower priors and therefore stronger compression. We used 9 approximately evenly spaced values of *p ∈* [1.10, 2.50] for the simulation, with the value of *p* for each panel indicated in the subtitle.

Figure 1 illustrates how capacity constraints (*C*_*i*_) and prior dispersion (*p*_*i*_) jointly shape the mapping from true value differences to internal representations. In each panel of Figure 1, for a channel *i*, we vary the capacity *C* while keeping the prior dispersion fixed: higher *C* leads to less compression, producing steeper encoding curves closer to the identity line. Across panels of Figure 1, we use the same set of the capacity *C* and vary prior dispersion *p*, showing that narrower priors (larger *p*, smaller Γ) yield stronger compression, while broader priors (smaller *p*, larger Γ) decrease compression. Narrower priors compress large differences because the prior mass is concentrated over small differences. Together, these simulations show that different degrees of nonlinearity in attribute value difference representations can arise from variations in channel capacity constraints, prior dispersion, or a combination of both factors.

### Predicted representations within a choice task

Now we move to the context of a multi-attribute choice in which the agent’s goal is to encode attribute value differences in a way that supports selecting the option with the higher overall subjective value.

The overall subjective value difference between two options is modeled as a weighted sum of the attribute-specific value differences:

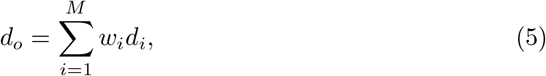

where *w*_*i*_ denotes the subjective weight assigned to attribute *i* (e.g., tastiness or healthfulness). As before in the case of an estimation task, the agent cannot directly access the true attribute differences *d* _*i*_, b ut i nstead r elies o n i nternal representations *r*_*i*_ ∼ *Q*(*r*_*i*_ | *d*_*i*_). The internally represented overall difference i s therefore

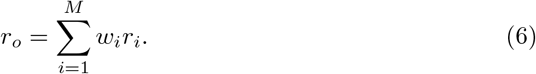

The agent is now optimizing not for estimation accuracy, but for decision performance: it seeks to encode the inputs such that the resulting choice probabilities are as close as possible to those that would arise from the true values. Formally, the objective is to minimize the *stakes-weighted cross-entropy* between the agent’s choice probabilities *p* (based on noisy internal signals) and the normative probabilities *q* (based on the true value differences). T he s takes, d efined as the absolute overall subjective value difference | *d*_*o*_|, weigh errors more heavily when choices are more consequential. The optimization is performed under a shared capacity budget *C* across all attribute channels enabling the objective discussed above. This shared budget allows capacity to be re-allocated across attribute channels according to their contribution to the downstream choice objective.

Taking the functional derivative of the corresponding Lagrangian and setting it to zero yields a generalized Gibbs-form encoder (see Methods subsection: “Lagrangian and decomposition” of “Choice task”):

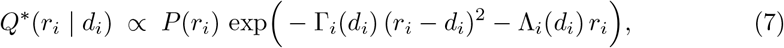

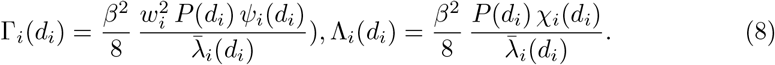

The first term, governed by Γ_*i*_(*d*_*i*_), determines the effective encoding precision, while the coupling coefficients *χ*_*i*_(*d*_*i*_) in the second term, Λ_*i*_(*d*_*i*_), capture how potential systematic biases in other attribute channels influence the encoding of channel *i*. If it is not zero, Λ_*i*_(*d*_*i*_) introduces a systematic shift in the encoded mean. In our experimental setting, however, these effects are expected to be negligible (see the sub-section Simplified regime and interpretation in the Methods for details). Therefore, we set *χ*_*i*_(*d*_*i*_) equal to zero, thus negating the Λ_*i*_(*d*_*i*_) term in the simulations in Figure 2.

**Fig. 2.**
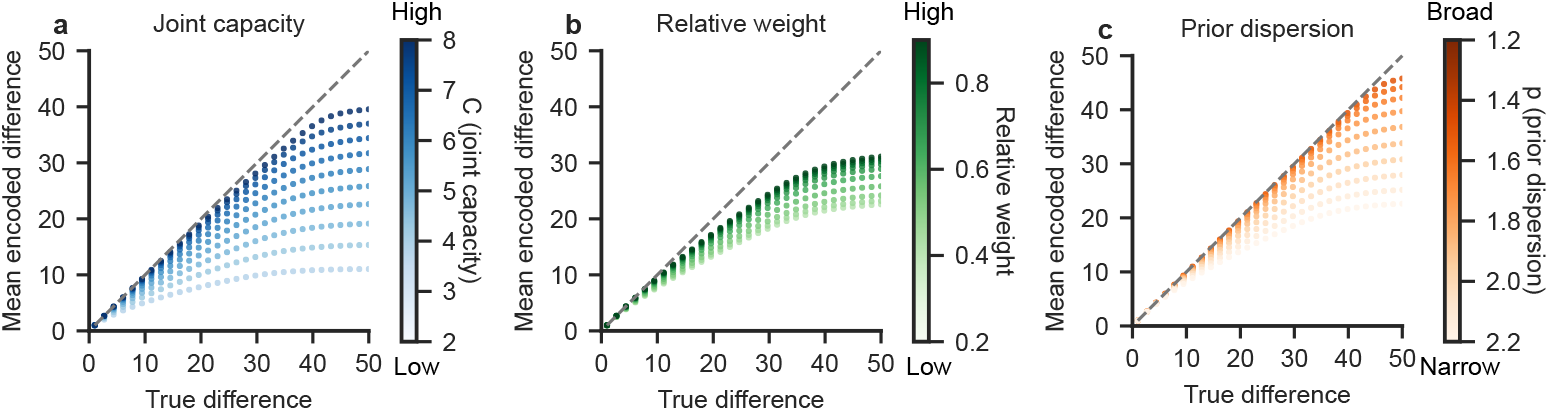
Joint channel capacity, subjective weight and prior dispersion modulate compression in choice task. Simulations for channel *i* follow Equation 36. The mean encoded difference responses 𝔼 [*r*_*i*_ | *d*_*i*_] is plotted as a function of the true difference *d*_*i*_. For the simulations, we fix *ψ*(*d*) = 1, *β* = 1. (a) We vary *C* (joint channel capacity) while fixing *w* = 1. As capacity *C* decreases, compression becomes stronger. We used 10 evenly spaced values of *C* ∈ [2, 8] to simulate. (b) We fix *C* = 5 and vary the relative subjective weight *w*. As *w* decreases, compression becomes stronger. We used 10 evenly spaced values of *w*_*i*_ ∈ [0.2, 0.9] to simulate. (c) The prior over encoded values was *P* (*d*) ∝ (1 + |*d*|^*p*^)^*−*1^. Prior dispersion was varied by sweeping the exponent *p* ∈ [1.2, 2.2], with channel capacity *C* = 5, subjective weight *w* = 1.0.

This results in the following reduced formulation:

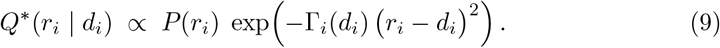

*ψ*_*i*_(*d*_*i*_) is the expected stakes given *d*_*i*_ and *β* captures decision noise at the level of the softmax choice rule. 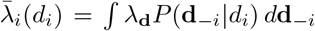 is the effective per-channel capacity multiplier obtained by marginalization (see Methods). Again, the Γ_*i*_(*d*_*i*_) term as a whole determines the effective encoding precision.

This formulation highlights a key difference from the estimation task. While the precision in the estimation task depends only on the channel capacity (*C*_*i*_) and the prior (*P* (*d*_*i*_)), in the choice setting it is additionally modulated by the subjective weights *w*_*i*_ that determine each attribute’s contribution to overall value. Thus, even if two attributes share the same prior, differences in their subjective weights will produce different effective encoding precision, both through the direct 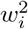 scaling in Γ_*i*_ and through the weight-driven reallocation of the shared capacity budget *C* across channels.

To illustrate these effects, Figure 2 shows simulations of the mean encoded difference responses under a simplified encoder where *ψ*(*d*) = 1 . In Figure 2a, we vary the channel capacity parameter *C* while keeping the weight fixed (*w* = 1): higher *C* again yields steeper, less-compressed curves. In Figure 2b, we fix *C* = 5 and vary the subjective weight *w*, showing that stronger weight increases effective encoding precision, producing less compression. In Figure 2c, we vary the prior dispersion: broader prior again yields steeper, less-compressed curves. This demonstrates that different degrees of nonlinearities in different attribute value differences in choice task reflect not only different capacity constraint or priors, but also differences in subjective weighting across attributes. In Supplementary Figures S10 and S11, we show that alternative forms where stakes increase or decrease linearly with |*d*| yield qualitatively similar results.

In the following sections, we test whether data from human participants making difference estimations and multi-attribute choices display the characteristics predicted by the theoretical account. Specifically, we test for evidence of the non-linear attribute difference representations during estimations and choices predicted by the power-law relationships in the model. The estimation task design allows us to ***exogenously control the reward for representing the difference*** in a given attribute accurately. The choice tasks allow us to investigate whether ***subjective weights are associated with systematic, attribute-specific encoding precision*** during multi-attribute choice.

### The degree of compression and variability in attribute difference estimations

We used an estimation task to determine if behavioral signatures of attribute encoding align with the model’s predictions. In this task, participants were required to report the difference in tastiness or healthfulness between two food items presented as pictures on a computer screen. Participants were informed that they needed to estimate the value differences in each attribute accurately in order to receive the highest payoff. We take the true attribute value differences to be those we calculated from the independent tastiness and healthfulness ratings provided for each food item when it was shown in isolation during the first phase of the task. Each food was rated in isolation twice and we used the average of both ratings as the true rating for a food item.

We began by testing if the relationship between reported estimates of the attribute value differences and true attribute value differences followed a power-law relationship as predicted. We modeled the relationship between estimated attribute differences and true differences (Fig. 3a) using either linear or power functions through hierarchical Bayesian modeling (Eq. 37 and Eq. 38, details in Methods section). As shown in model comparison Table 1 and Table 2, across both tastiness and healthfulness, the power function consistently provided a better fit compared to the linear model. The tastiness attribute dimension showed more nonlinearity/compression than the healthfulness attribute dimension (Figure, 3b, 95 % High Density Intervals (HDI) for tastiness’ exponent = [0.566, 0.718], healthfulness = [0.862, 1.287]). Note that within the estimation task alone, we cannot definitively distinguish whether the compression difference between attributes reflects different channel capacities *C*_*i*_, different prior dispersions *p*_*i*_, or a combination of both. The model predicts the same qualitative pattern under either account.

**Table 1.** Model comparison for tastiness difference estimation.

|  | elpd_diff | se_diff |
| --- | --- | --- |
| **power-law** | 0.0 | 0.0 |
| **linear** | -48.1 | 10.0 |

**Table 2.** Model comparison for healthfulness difference estimation.

|  | elpd_diff | se_diff |
| --- | --- | --- |
| **power-law** | 0.0 | 0.0 |
| **linear** | -42.7 | 10.8 |

**Fig. 3.**
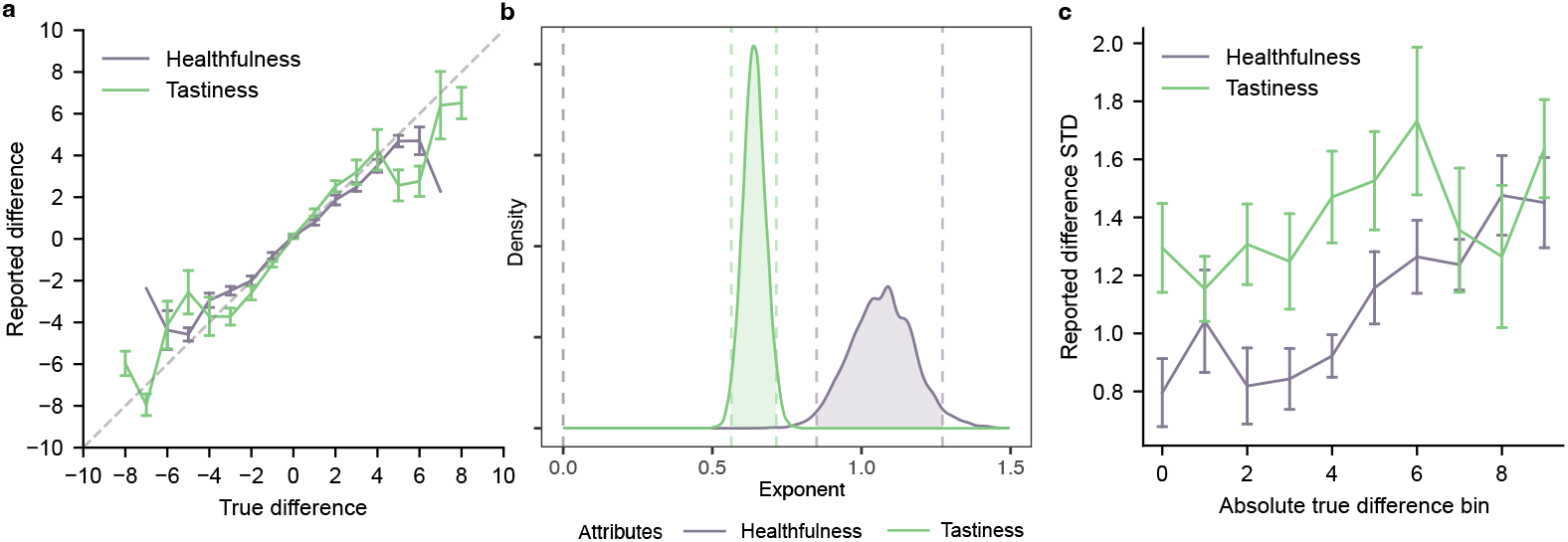
Attribute-specific nonlinear compression and variance in attribute difference estimation. (a) Relationship between true attribute differences and reported differences in the estimation task for tastiness (green) and healthfulness (purple). Points show binned means with error bars indicating s.e.m. (b) Posterior distributions over the power-law exponent from the hierarchical Bayesian power model for tastiness and healthfulness, showing stronger compression (lower exponent) for tastiness. Shaded regions indicate 95% HDIs. (c) Standard deviation of reported differences as a function of 10 quantile bins of absolute true difference, showing greater variability for tastiness than for healthfulness. Points show binned means with error bars indicating s.e.m.

The difference in compression between the tastiness and healthfulness dimensions allows us to assess a key model signature. Specifically, the model predicts a dual signature of resource limitation: both channel capacity, *C*_*i*_, and the prior dispersion, *p*, shape not only the mean but also the variance of the posterior distribution *Q*(*r*|*d*). This link between bias and variability, where compression and uncertainty emerge jointly from resource-limited encoding is confirmed by model simulations (Supplementary Figure S4): for realistic, sufficiently high capacities, posterior variance decreases monotonically with capacity; and it increases with broader priors. Empirically, we observed a pattern in the estimation task consistent with the model simulations: tastiness, which showed stronger compression, also exhibited higher variance than healthfulness when estimating attribute differences (Figure 3c). A linear mixed-effects model allowing different variances per attribute (Equation 39) fits the data significantly better than a homoscedastic model (Supplementary Table S8), with the estimated residual standard deviation for difference in healthfulness ratings being 0.771 times that of tastiness ratings.

### Choices are based on nonlinear transformations of attribute value differences

The goal-adaptive difference encoding model yields two empirically testable signatures in human multi-attribute choice behavior. First, as in estimation, choices reflect *nonlinear transformations of attribute differences*. Second, the degree of nonlinearity expressed for each attribute depends not only on environmental structure and total capacity, but also on the attribute’s subjective importance for the decision-maker’s goals. While these signatures are not uniquely diagnostic of the resource rational mechanism, their presence provides targeted evidence that the choice process departs from linear, WAD-style integration in the manner the theoretical account anticipates.

Unlike the estimation task, participants’ responses in a choice task do not directly indicate their perception of the attribute differences. We must utilize a model of the decision process to estimate the functional form of the relationships between attribute differences and binary decisions. We used the drift diffusion model for this purpose because it is an established process-level model linking latent value signals to both choices and response times. The use of both choice and response-time data allows for more accurate parameter estimation relative to softmax functions that rely on choice data alone [26, 27]. We specified and fit both linear (baseline) and nonlinear choice models, which assumed that choices are made based on the sum of the weighted linear or nonlinear differences in each attribute, respectively. The nonlinear choice models included power-functions applied to the attribute value differences. Power functions offer a flexible approach to modeling nonlinear transformations: they encompass the linear case (when the exponent is 1) and can produce convex or concave curvature depending on the parameterization. As a complementary approach to ensure that our results are not dependent on the specifics of the power functions used within the drift diffusion model, we also used segmented logistic regression, which does not impose a fixed functional form on the decision process. These additional analyses are reported in the Supplementary Information.

We used hierarchical Bayesian modeling to fit the linear and nonlinear difference models to multi-attribute decisions about food and monetary distributions between oneself and another person (Dictator games). In total, we test four separate multi-attribute choice datasets across the two domains. For clarity and conciseness, we generally report only the results from one food choice dataset (Food1) and one dictator game dataset (Social1) in the main text and include the two conceptual replications (Food2, Social2) in the supplementary materials.

The model comparisons were performed using approximate leave-one-out cross-validation [28] (details in Methods - “Bayesian hierarchical model comparison via leave-one-out cross-validation (LOO)” section). Table 3, Table 4 and Supplementary Table S2, S3 and S5 show model comparison results across the different tasks and conditions, evaluated using the expected log pointwise predictive density (ELPD). The simulations based on both models are shown in Figure 4, Figure S5 and Figure S6.

**Table 3.**
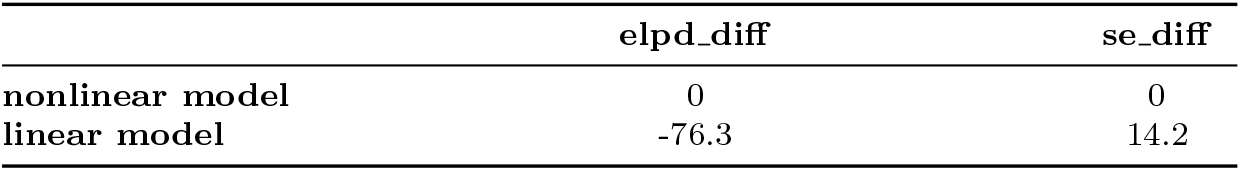
Model comparison for the “Images” condition in Food1 dataset.

|  | **elpd_diff** | **se_diff** |
| --- | --- | --- |
| **nonlinear model** | 0 | 0 |
| **linear model** | -76.3 | 14.2 |

**Table 4.**
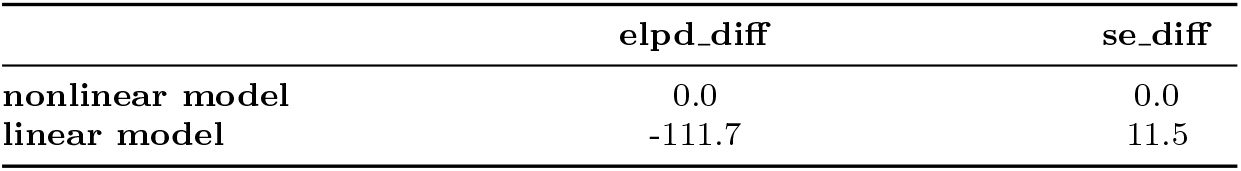
Model comparison for the “Time-free” condition in Social1 dataset.

**Fig. 4.**
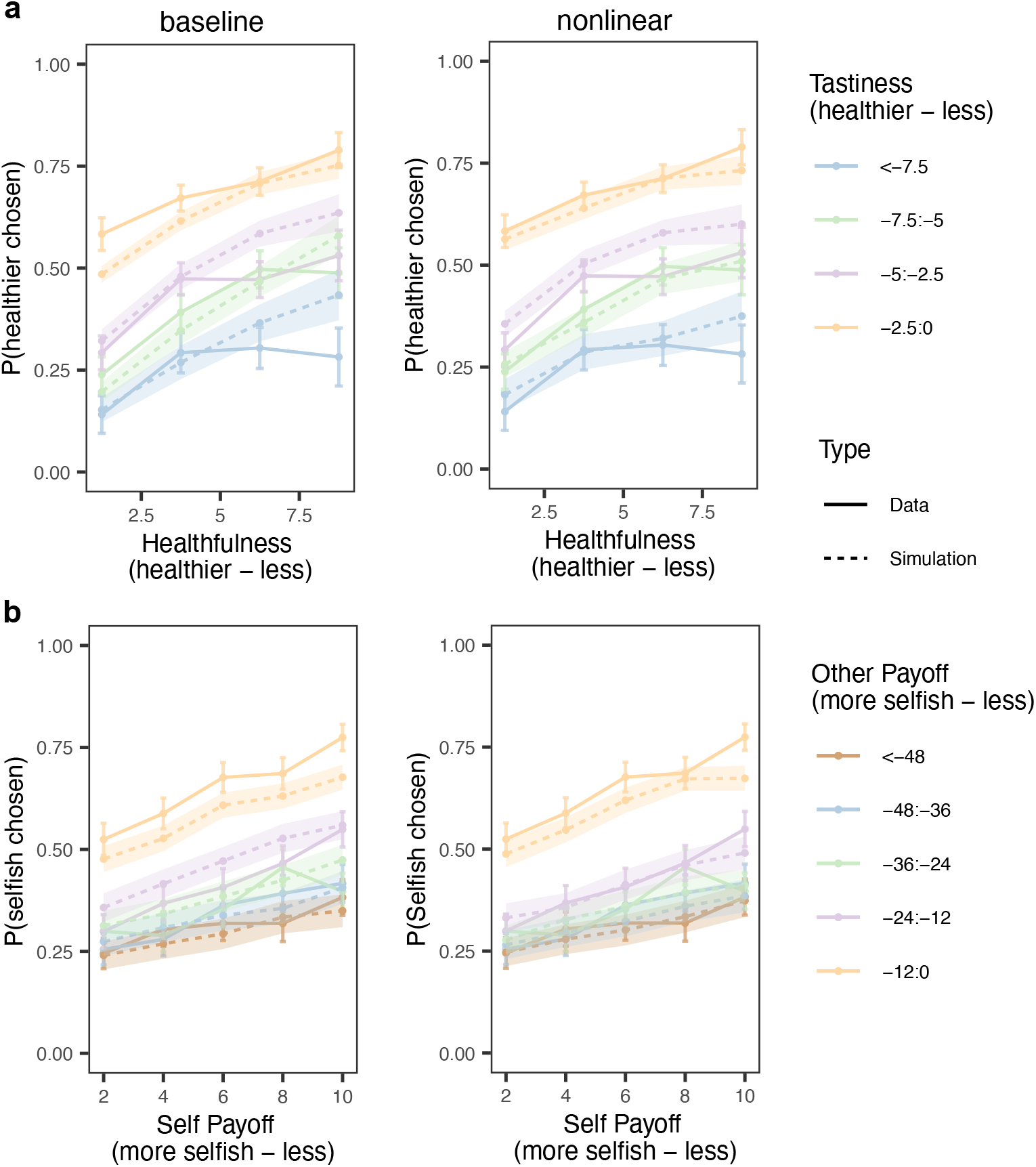
Nonlinear difference representations better capture multi-attribute choice in food and social decisions. (a) Food choices (Images condition in Food1 dataset). The y-axis shows the probability of choosing the healthier option as a function of the healthfulness difference (x-axis; healthier option minus less healthier option) between options, shown separately for bins of tastiness difference (healthier option minus less healthier option; color coded). Solid lines denote empirical data, and dashed lines denote simulations based on the choice models. Left panels show simulations from the baseline model that assumes linear differences and right panels show simulations from the model incorporating nonlinear differences. (b) Dictator game choices (Time-free condtion of Social1 dataset). The y-axis shows the probability of choosing the more selfish option as a function of self-payoff differences (x-axis; more selfish option minus less selfish option), plotted across bins of other-payoff differences (more selfish option minus less selfish option; color coded). Across both domains, the nonlinear difference models more accurately capture the curvature in choice patterns.

These results indicate that the choice model allowing for a power-function transformation of the attribute differences yields a better fit than the baseline choice model using linear differences. *This finding supports the resource-rational encoding model prediction that choices are based on nonlinearly encoded attribute differences*.

### The degree of nonlinear encoding is proportional to an attribute’s subjective importance

The resource-rational difference encoding model predicts that the degree of nonlinearity in attribute difference encoding is proportional to an attribute’s subjective importance to the decision-maker’s goals. To evaluate this implication in choices, we compared the fitted exponents of the power functions across different attributes. Eye-tracking data from participants in the Food1 dataset performing an analogous version of the food choice task that showed only their previous attribute ratings instead of pictures of the foods themselves, indicate that they allocate more attention resources to tastiness than healthfulness attributes during subjective choices. Specifically, the participants spent more time fixating on tastiness relative to healthfulness (Equation 40; intercept (difference) = 176.00 ± 71.19, *p* = 0.0147) and a greater proportion of fixation on tastiness relative to healthfulness (Equation 41; *β* = 0.32 ± 0.12, *p* = 0.0080), suggesting that more attention and subjective weight was given to tastiness attributes during food choices. Importantly, subjective attribute weights do not influence the estimation task, because rewards are solely determined by the accuracy of estimating attribute differences, with all attributes assigned equal importance exogenously.

Recall that, in the estimation task, tastiness ratings were more variable and encoded more nonlinearly than healthfulness attributes. In the choice task, if the subjective weight on tastiness is sufficiently higher than healthfulness attributes, as is indicated by the fixation data, then the resource-rational encoding model predicts that the relative magnitude of the exponents can reverse in the choice compared to ratings experiments. In other words, the exponent on tastiness differences will be closer to 1 (i.e., linear) than that on healthfulness.

Indeed, we find that, in contrast to the estimation task results, the encoding of tastiness changes to become more linear than healthfulness during subjective choices. In both food-choice datasets, the exponent for the tastiness attribute was greater than the exponent for healthfulness attribute and was closer to 1 (leading to a more linear representation). Specifically, in Food1 dataset, the 95 % HDI of the fitted population-level exponents for tastiness and healthfulness were [0.84, 1.05] and [0.36, 0.62], respectively (Figure 5a). The 95 % HDI of the difference (tastiness - healthfulness) in exponents was [0.285, 0.624]. The Food2 dataset showed a similar pattern: the 95 % HDI for difference in exponent is [0.0968, 0.5112], with 95 % HDIs of [0.82, 1.21] for tastiness’ exponent and [0.61, 0.80] for healthfulness’ exponent (Figure S8). As shown in Figure 5b and Figure S8, across both food-choice datasets, the majority of participants (85 % - 87 % in Food1 dataset, 88 % in Food2 dataset) exhibited higher subject-level median exponents for the tastiness attribute compared to the healthfulness attribute.

**Fig. 5.**
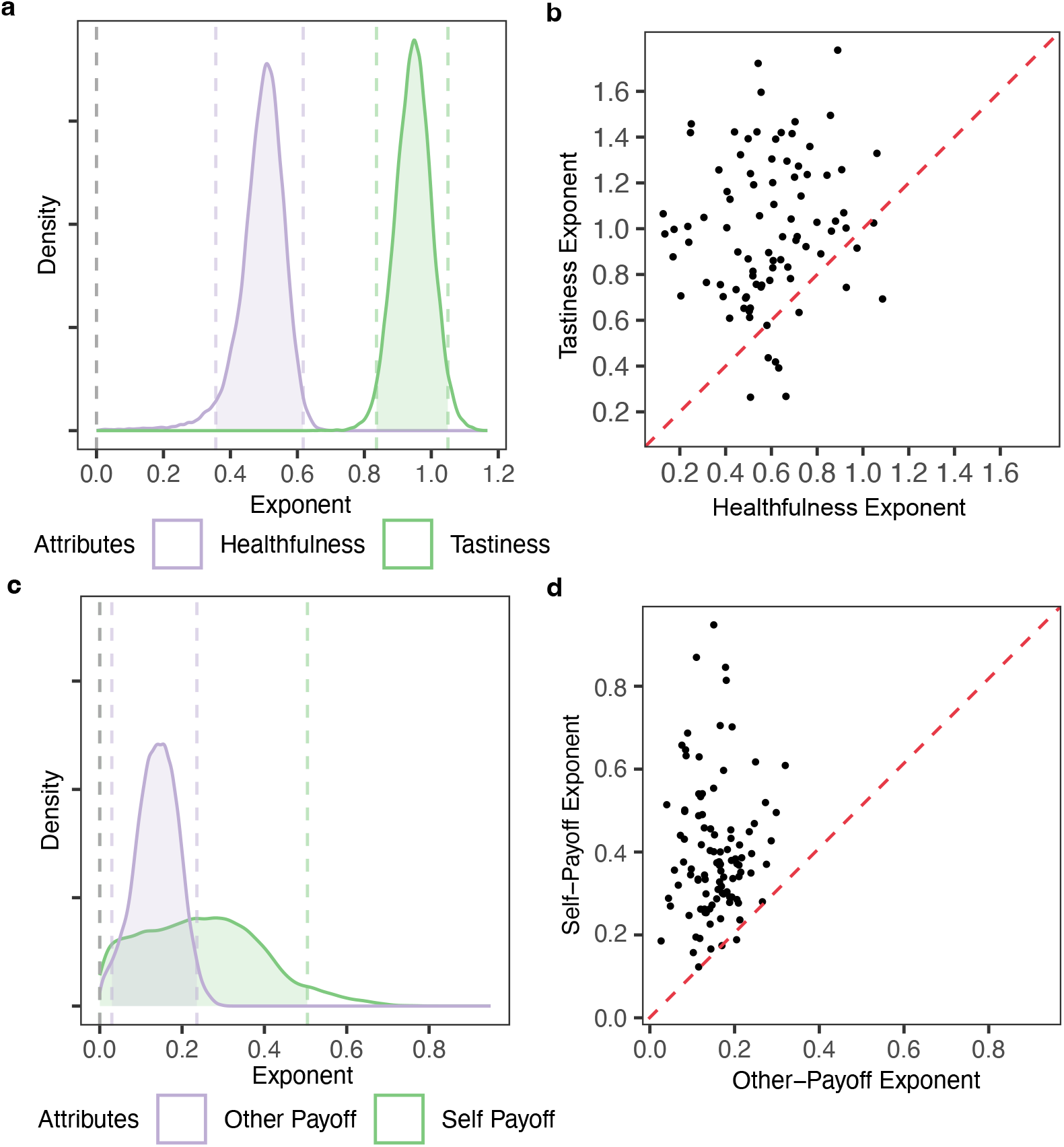
Attribute-specific nonlinearity across decision domains. (a) This density plot shows the population-level posterior distributions of exponents for tastiness and healthfulness in the Food1 dataset. Tastiness shows a higher exponent, closer to linear encoding, than healthfulness. Shaded areas indicate 95% HDIs. (b) This scatter plot shows subject-level posterior median values for the exponents for tastiness versus healthfulness differences in Food1. Most participants lie above the unity line (red), indicating stronger (more linear) encoding of tastiness than healthfulness. (c) This density plot shows the population-level posterior distributions of exponents for self-payoff and other-payoff attributes in the Social1 dataset, revealing substantial inter-individual variability and generally stronger encoding of self-payoffs. (d) This scatter plot shows subject-level posterior median values for the exponents on self-payoff versus other-payoff differences, with the majority of participants exhibiting higher exponents for the self-payoff attribute.

Social choices also showed asymmetry in the degree of attribute nonlinearity, although with more individual variability in exponent values than food choices. This may reflect the high degree of variability in social preferences across participants. Overall, in the free-response condition, the majority of participants (97% in Social1 dataset, 80 % in Social2 dataset) exhibited higher subject-level posterior median values for the exponent on own-payoff attribute compared to the others-payoff attribute (Figure 5d and Figure S9). This asymmetry in exponent values is consistent with an encoding scheme that depends on an attribute’s subjective importance and the fact that the exponents on self-payoffs are larger is in line the with the general finding from many studies using the dictator game that people, on average, place more importance on their own payoffs than payoffs for others [29]. Additionally, the overall degree of nonlinearity in payoff representations, especially for self, varied substantially across participants (Social1 population-level self payoff 95% HDI = [3.9 × 10^−6^, 0.50], other payoff = [0.029, 0.24]; Figure 5c; Supplementary Figure S9). The inter-individual variability in exponent magnitudes - our proxy for encoding precision - may be driven, at least partially, by differences in the degree of resources participants have or make available for the decision task.

### Increased resource constraints lead to a more nonlinear encoding

Manipulating time pressure is a commonly used method of experimentally varying resource constraints [30–32]. As shown in Figure 2, the theoretical model predicts that increasing the capacity constraint (lower *C*) produces stronger compression of value differences in the choice task. Consistent with the model, the fitted exponents of subjective value differences were generally smaller (more compressed) under time pressure than that in time-free conditions across both “Social 1” and “Social 2” datasets, indicating that increased constraint leads to greater compression of subjective representations (Figure 6; parameter recovery shown in Figure S17 and Figure S20). Quantitatively, the 95% HDIs for the differences between time conditions were as follows: “Social 1”—self: [−0.241, 0.547] with 78.7% posterior mass *>* 0; other: [−0.101, 0.212] with 81.3% posterior mass *>* 0; “Social 2”—self: [−0.674, 0.625] with 63.3% posterior mass *>* 0; other: [0.146, 0.366] with 100% posterior mass *>* 0. Note that the relatively high proportion of posterior mass near the lower bound of zero in the time-free conditions limits the degree to which time-pressure can cause further compression. Accordingly, we observe the most substantial change in posterior mass for other payoffs in the Social 2 data set, which has the least posterior mass at the lower bound (0) in the time-free trials. These results provide empirical support that manipulating available processing time modulates representational capacity in the manner predicted by the model. Consistent with prior work, time pressure does not simply increase random noise; instead, decision-makers adapt their information-processing strategies to optimally allocate limited cognitive resources under external computational constraints [32]. While we acknowledge that increasing time pressure may have additional effects on cognition and behavior that are not mediated by resource constraints, these results are nevertheless consistent with the predictions of the goal-adaptive difference encoding model.

**Fig. 6.**
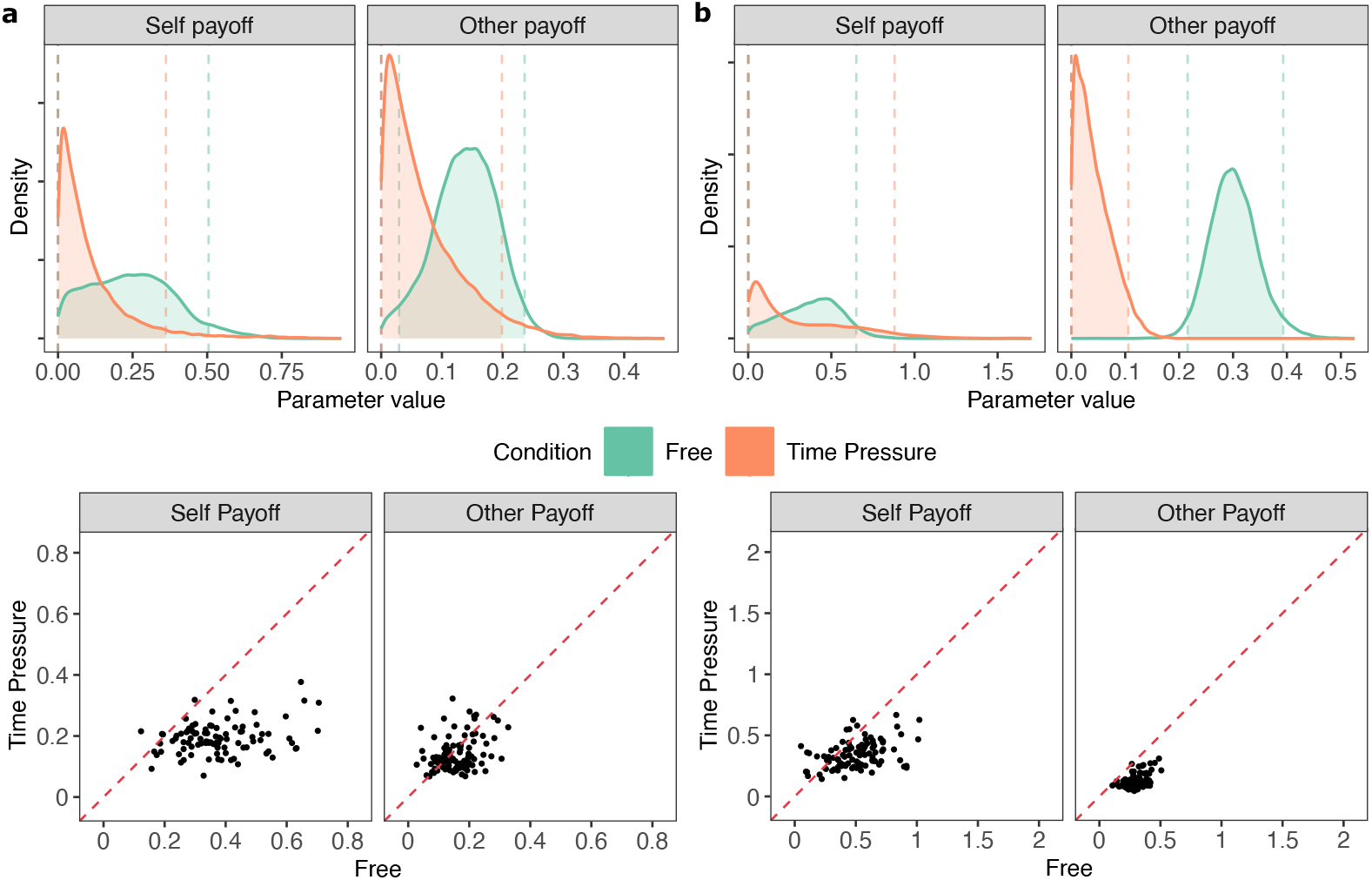
Time pressure increases compression of subjective value differences in social choice. The upper panel shows posterior distributions of the power-function exponents governing nonlinear encoding of attribute differences separately for self-payoff and other-payoff attributes and for time-free versus time-pressure conditions. Exponents are reduced under time pressure (i.e., further from 1), indicating stronger compression of subjective value differences when available processing time is constrained. Shaded areas indicate 95% HDIs. The lower panel shows subject-level posterior median values for the exponents fitted in each condition. Most participants lie below the unity line, indicating stronger (more linear) encoding in Time-free condition than in Time-pressure condition. (a) Results for Social1 dataset. (b) Results for Social2 dataset.

## Discussion

We propose a theoretical account in which resource-rational compressed representations of attribute value differences drawn from emergent decaying priors, producing compressed representations that are further shaped by task goals. We show that this compression can lead to choice behaviors better described by nonlinear attribute value differences, deviating from the predictions of a baseline linear-difference choice model and other WAD-based models. Our results provide evidence that, in estimation, attribute-specific variation in nonlinearity is compatible with differences in capacity constraints and/or the dispersion of prior beliefs. In choice, the same framework includes an additional role for subjective attribute weighting, which can redistribute encoding resources across attribute channels and thereby modulate the nonlinearity expressed during decision making. Together, these findings offer a coherent account of why the relative distortions in attribute differences can vary across tasks.

Our model builds on a core principle shared with efficient coding theories—that neural processing is shaped by the statistical structure of the environment [33, 34]—and with rate-distortion theory, which posits that performance is optimized under constraints on information-processing capacity [35]. Both frameworks have been successfully applied to explain a range of perceptual phenomena, including perceptual biases [36–38], numerosity estimation and discrimination [13, 39–42], limitations in visual working memory [43, 44], and perceptual generalization [45]. Our attribute value estimation task and multi-attribute choice task share key features with these paradigms: participants perceive stimuli and either estimate particular aspects or discriminate between alternatives. This alignment makes it natural to extend efficient coding and rate-distortion principles to the domain of multi-attribute valuation and decision-making.

Nevertheless, our value estimation and multi-attribute choice tasks differ from perceptual and numerosity tasks in that they likely involve higher-level goal-selection and reward-related processes. A growing body of work suggests that decision-makers in risky choice lack direct access to objective expected payoffs and instead rely on capacity-constrained internal representations [46–49]. Moreover, individual differences in risk aversion have been linked to the precision of neural representations of numerical magnitude [50], contrasting with classical accounts that attribute risk aversion to later-stage reward processing [51, 52]. Relatedly, efficient-coding accounts of subjective value can explain uni-dimensional valuation and choice [1]. Together, these findings align with our proposed framework for multi-attribute choices more generally: rather than explaining nonlinearity through higher-level processes, we argue that it originates in the fundamental representations of differences, which are tuned by goals. Consistent with this view, sensory coding has been shown to adapt to behavioral goals [53–55], and functional remapping of stimulus–reward contingencies in early sensory cortices have been found to depend causally on top-down control signals from prefrontal regions [55, 56].

Our resource-rational model is formulated over attribute-value differences, treating individual attribute values as upstream inputs that may or may not already be distorted. This assumption that attribute differences are represented is supported by extensive evidence indicating that both humans and monkeys frequently engage in attribute-wise comparisons during multi-attribute decision-making, as shown through eye-tracking, mouse-clicking, and computational modeling studies [57–60]. Additionally, various neural-level findings (fMRI or neural recording) have provided support for within-attribute comparisons [61] and representations of value-difference signals [62, 63]. Importantly, our account does not preclude resource-rational encoding of individual attribute values in addition to their differences. Indeed, standard economic assumptions such as diminishing marginal utility already imply systematic nonlinearities at the level of single-attribute values. Critically, encoding single values and then subtracting them is not generally mathematically equivalent to encoding differences directly: upstream nonlinear value encoding can induce difference signals that depend on absolute levels as well as on differences, whereas direct difference encoding predicts invariance to common-mode shifts that leave differences unchanged. Determining how distortions are distributed across these two representational levels—only at the individual-value level, only at the difference level, or at both levels—will require more targeted tests. Nevertheless, assuming at minimum that difference signals are encoded—as suggested by the converging evidence above—our proposal provides a principled account of how systematic distortions can arise in the internal representation used for multi-attribute comparison.

Additionally, our proposal of resource-rational transformation of attribute-value differences is not the only indication in the literature that models based on non-linear differences can best account for multi-attribute choice behavior. Related prior work has repeatedly found that nonlinear difference-based models provide superior fits across diverse datasets [64–67]. However, previous studies have not provided a mechanistic explanation for why the transformation should be nonlinear. Our work explaining such effects within a resource-rational framework may generalize to these settings and provide an mechanistic account of the observed nonlinearity.

Our empirical tests provide several results in support of the proposed resource-rational difference encoding account. The multiple behavioral signatures we document form a coherent pattern across estimation and choice tasks, although they do not uniquely identify each component of the model. In particular, similar behavioral patterns can be generated by different combinations of capacity constraints, prior dispersion, and goal-dependent resource allocation, and our current tests cannot fully disentangle these contributions in isolation. Importantly, the time-manipulation condition in social choice provides a first step toward more direct tests of resource limitation, but designing experiments that cleanly target the remaining components is nontrivial. For example, manipulating subjective importance via explicit weight ratios in incentivization may prompt participants to adopt a different choice strategy (e.g., explicit arithmetic integration of weights and attribute values), altering the decision rule rather than selectively modulating resource allocation. Future work will therefore need to develop designs that isolate these components more selectively while guarding against such strategic and interpretational confounds. Fortunately, common choice analysis frameworks (e.g., logistic regressions and sequential sampling models) can be modified readily to include power-law functions that allow for compressed difference representations without requiring direct identification of capacity, priors, or goals. As we have shown here, such models better explain in-sample behavioural patterns and (expected) out-of-sample performance. Choice models augmented with power-law functions can be used to test the predicted influences of changes in priors, channel capacity, or goals.

Importantly, our findings do not necessarily suggest that nonlinearities in multi-attribute choice behavior stem solely from resource-rational representations of attribute value differences. Numerous theories have proposed additional or alternative mechanisms, each relaxing one or more of the core assumptions of the WAD framework to better capture empirically observed behaviors. For instance, some models introduce mechanisms that amplify or reduce the perceived similarity between options during comparison, employing various psychological mechanisms, including loss aversion [8], emotions [9], and attention [68, 69]. Another influential class of models, inspired by biophysical constraints on neural representation, suggests that the brain evaluates options using various forms of normalization among options [70, 71]. Furthermore, models incorporating inhibitory competition have been proposed to account for interactions between options and attributes [72, 73]. In parallel, attribute weights can dynamically vary throughout the decision process, modulated by shifts in attention [59]. That being said, our model provides a parsimonious account that simultaneously explains behavioral patterns observed in both the attribute value difference estimation task and the multi-attribute choice task, as the representation of value difference is essential in both tasks. It highlights that imprecision in attribute value representations is one of multiple mechanisms contributing to individual patterns in multi-attribute decision-making.

Our findings have important implications for real-world multi-attribute decision-making. Given that value differences in lower-weighted attributes are more compressed, promoting choices based on those attributes (e.g., healthfulness) may be difficult to accomplish by merely offering options that are more extreme on that dimension (e.g., even healthier), because at the level of internal representations the incremental advantage is more strongly compressed for large differences. In other words, the perceived advantage of a more healthful food reaches a plateau and does not continue to increase as the objective advantage grows. Instead, approaches that increase the subjective weight placed on healthfulness—by shaping beliefs, goals, or attention [74] toward health-related information—may be more effective. Similarly, factors that increase or decrease the total capacity across all channels such as sleep quality, stress levels, or distraction should also lead to corresponding decreases or increases in attribute difference compression and more or less constrained-optimal choices across domains.

## Methods

### Goal-adaptive difference encoding model

#### Estimation task

In the estimation task, participants were asked to report the difference in a single attribute between two options — such as the difference in tastiness or healthfulness between two food options. Let *d*_*i*_ denote the true value difference in attribute *i*. Individuals do not have direct access to *d*_*i*_, but instead form an approximate internal representation *r*_*i*_ with cognitive and sensory limitations. We assume it is drawn from a conditional encoding distribution *r*_*i*_ *∼ Q*(*r*_*i*_ | *d*_*i*_).

##### Formal objective

The agent’s goal in the estimation task is to report value differences as accurately as possible. This corresponds to a loss function of minimizing expected squared error between the internal representation and the true value difference, as shown in equation 1:

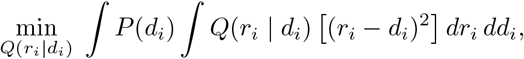

where *P* (*d*_*i*_) is the environmental distribution of the true value differences for attribute *i*.

##### Constraints

However, the encoding process is constrained by limited information-processing resources. For each attribute channel and each encoding instance, this constraint on information-processing capacity can then be formalized as a bound *C*_*i*_ on the Kullback–Leibler (KL) divergence [22] between the encoding distribution *Q*(*r*_*i*_ | *d*_*i*_) and the prior *P* (*r*_*i*_), where the prior captures the environmental expectations of the distribution of value differences for attribute *i*, as shown in Equation 2:

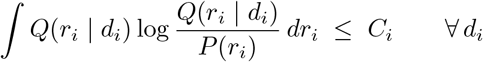

Additionally, the encoding distribution must be normalized:

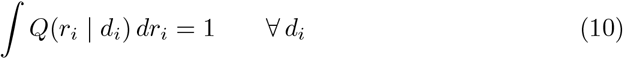

##### Lagrangian and decomposition

To solve this constrained optimization problem, we introduce a Lagrangian that combines the squared-error objective with the information-processing capacity and normalization constraints. The constraints are weighted by Lagrange multipliers 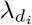 and 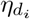 for the KL divergence and normalization constraints respectively, which quantify the tradeoffs between representational accuracy and information cost:

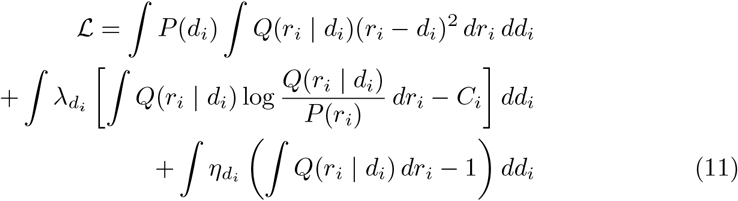

Taking the functional derivative with respect to *Q*(*r*_*i*_ | *d*_*i*_) and setting it to zero yields the equation:

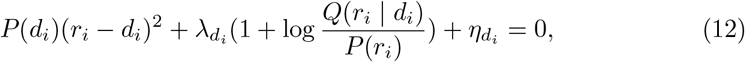

which gives us the optimal encoding distribution, shown as Equation 3:

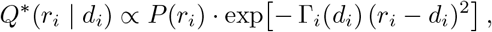

where 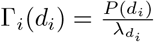 denotes the effective encoding precision for attribute channel *i* at value difference *d*_*i*_. Note that *C*_*i*_ is constant across all values of *d*_*i*_ for a given channel.

#### Choice task

In the choice task, participants made choices between two options that differed along multiple attributes, such as tastiness and healthfulness. The goal of this task is to choose the option with the higher subjective value.

We assume that participants make decisions based on the subjective difference in value between the two options. Let *d*_*i*_ denote the true difference in attribute *i* between two options in a choice set. The true overall subjective value difference between the two options is then defined as Equation 5:

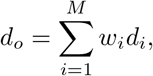

where the weights *w*_*i*_ reflect the participant’s subjective importance over attribute *i*.

Due to information-processing limitations, the agent cannot access the true values *d*_*i*_ directly. Instead, each attribute difference is encoded through an independent noisy channel *r*_*i*_ ∼ *Q*(*r*_*i*_ | *d*_*i*_), and the agent computes an internal estimate of the overall subjective value difference, shown as Equation 6:

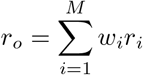

##### Formal objective

The agent’s goal is to encode value differences between options in a way that supports making the choice with higher value. Formally, the objective is to minimize the *stakes-weighted cross entropy* between the agent’s choice probabilities *p*, which depend on noisy internal representations of the value differences, and the normative target probabilities *q*, which are derived directly from the true value differences. Here, the *stakes* are defined as the absolute overall value difference between the two options, |*d*_*o*_|, which quantifies how costly it is to make a mistake: errors on trials with large value differences matter more than errors on small ones.

Because the attribute channels are assumed conditionally independent given their respective value differences, the joint encoding distribution factorizes as

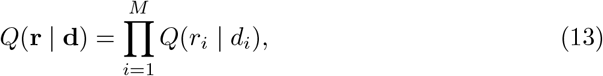

where **d** and **r** are the vectors for true and represented attribute value differences across attributes.

The encoders 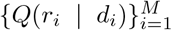 are constructed to minimize the *stakes-weighted* expected cross entropy between the normative target choice probability *q* and the choice probability distribution *p* that follows from the agent’s noisy internal representations:

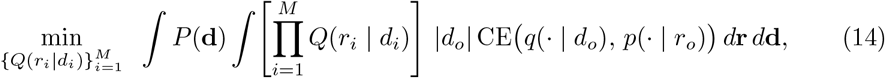

where the cross entropy is

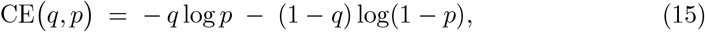

and *P* (**d**) is the environmental distribution of the true value differences.

##### Constraints

The total information processing across all attribute channels is limited, where *C* is the capacity bound:

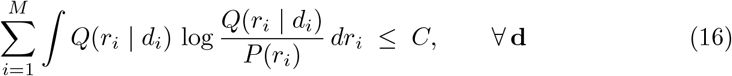

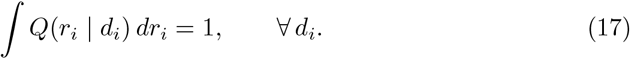

##### Quadratic upper bound for cross entropy

The cross-entropy between a target probability *q* ∈ (0, 1) and a model probability *p* ∈ (0, 1) is

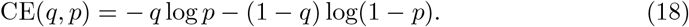

This is the standard loss function used to measure how well probabilistic choice policies match a normative target.

In discrete-choice models, the probability *p* is typically expressed as a smooth function of a weighted representation of the value difference. For example, consider the logistic choice rule through which an agent makes a choice *c* between two options {0,1}, in this case we have

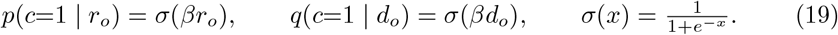

Here, *β* is the inverse temperature parameter controlling the amount of downstream decision noise.

The logistic cross-entropy loss

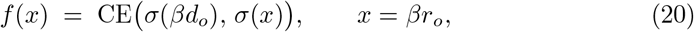

is convex with second derivative bounded by 1*/*4, which means it is an *L*-smooth function with smoothness constant 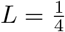 .

A general property of *L*-smooth functions [75, 76] is that they admit a quadratic upper bound around any point *x*^*^:

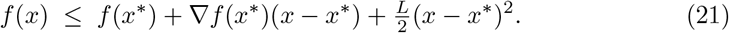

This inequality states that the first-order Taylor expansion of *f* at *x*^*^, plus a quadratic correction of curvature ^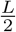^, always upper-bounds the function.

Specializing to our case, we expand around the target-matching point *x*^*^ = *βd*_*o*_, where the model prediction matches the target and thus ∇*f* (*x*^*^) = 0. The linear term vanishes, leaving only the quadratic correction:

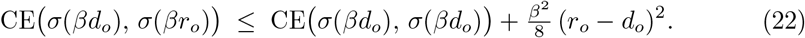

Here CE *σ*(*βd*_*o*_), *σ*(*βd*_*o*_) is the entropy of the target distribution. Since it depends only on *d*_*o*_ and not on the encoder, it is constant with respect to optimization and can be dropped in subsequent steps.

##### Lagrangian

Introducing Lagrange multipliers *λ*_**d**_ *>* 0 for the KL constraints. There is one multiplier per joint realization **d**, reflecting that the shared capacity budget must hold for every **d** separately, and 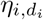 for the per-channel normalization constraints. The Lagrangian is:

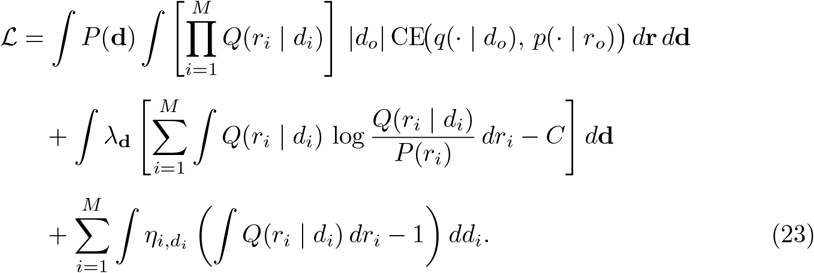

Plugging the quadratic upper bound into the choice objective and dropping the constant entropy term CE(*σ*(*βd*_*o*_), *σ*(*βd*_*o*_)), which depends only on *d*_*o*_ and not on the encoder, gives the simplified Lagrangian:

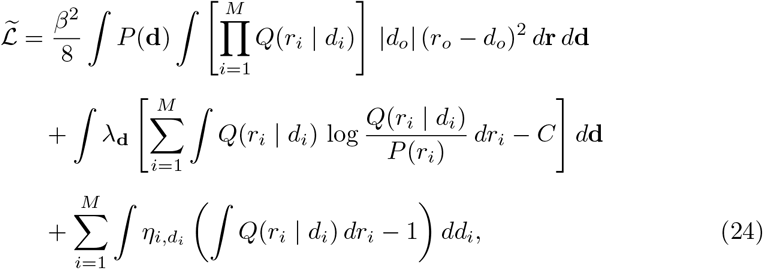

with *r*_*o*_ = ∑*i w*_*i*_*r*_*i*_ and *d*_*o*_ = ∑_*i*_ *w*_*i*_*d*_*i*_. The multiplier *λ*_**D**_ is shared across all attributes for a given **d**, allowing the optimal capacity allocation across attributes to emerge from the optimization rather than being fixed in advance. This expression makes clear that the effective loss combines a quadratic distortion penalty with information-processing costs.

##### Decomposition

The quadratic distortion term in the bounded Lagrangian is

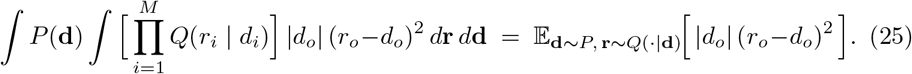

Expanding 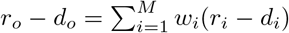 gives

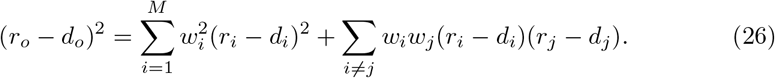

Taking the conditional expectation given **d**, the cross-terms involve expectations of the form

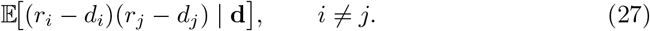

We can expand each difference around its conditional mean:

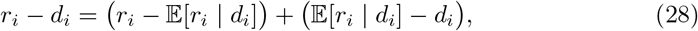

and similarly for *r*_*j*_ − *d*_*j*_. Expanding the product gives four terms:

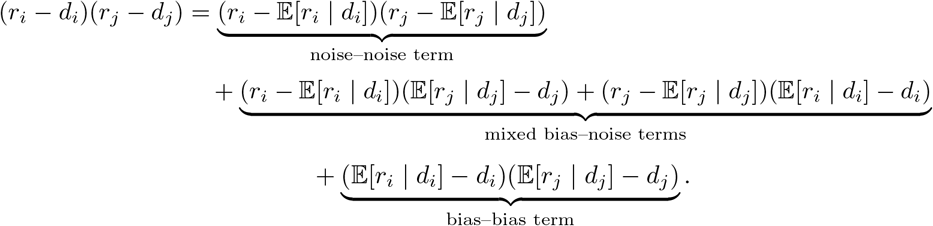

Taking the conditional expectation given **d**, the mixed terms vanish because each residual (*r*_*i*_ − 𝔼[*r*_*i*_ | *d*_*i*_]) has zero conditional mean:

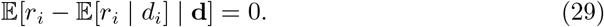

This leaves

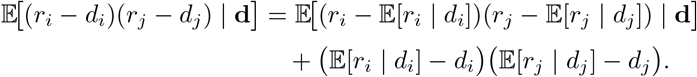

The first term captures any correlated encoding noise across channels, and the second term captures correlated bias across channels. Under the assumption of conditionally independent encoding noise, the first term vanishes. The remaining cross-term is thus fully determined by systematic biases, yielding

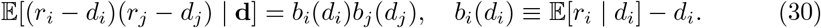

Consequently, the quadratic term decomposes into an independent channel-wise distortion term and a bias–bias coupling term:

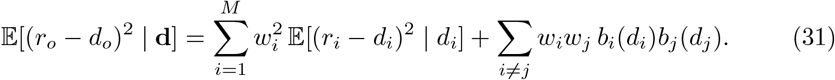

Taking the expectation over the environmental distribution and weighting by the stakes |*d*_*o*_| gives

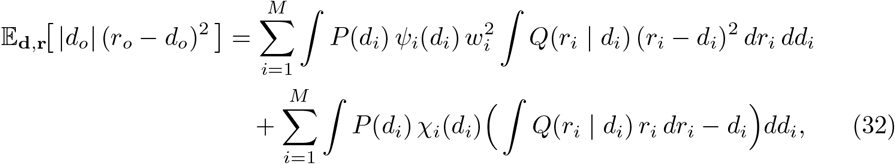

where

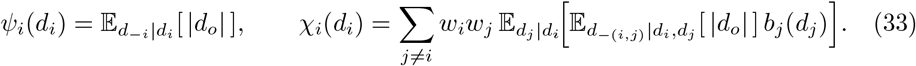

Here *ψ*_*i*_(*d*_*i*_) represents the expected stakes given *d*_*i*_, and *χ*_*i*_(*d*_*i*_) captures how biases in other channels influence encoding in channel *i*.

Substituting these components into the bounded optimization problem gives the full Lagrangian:

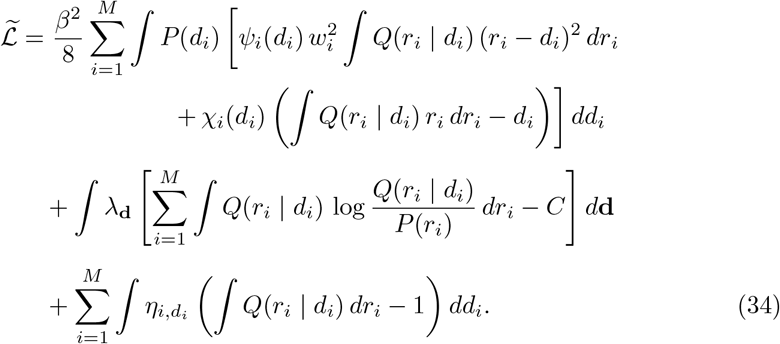

Taking the functional derivative 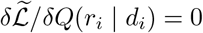, the constraint term naturally produces a per-channel effective multiplier by marginalizing *λ*_**d**_ over **d**_−*i*_:

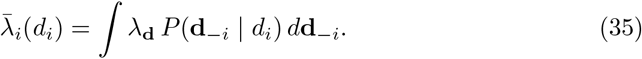

Solving for *Q*^*^(*r*_*i*_ | *d*_*i*_) yields a generalized Gibbs–form encoder, as shown in Equation 8:

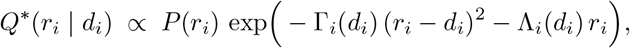

With

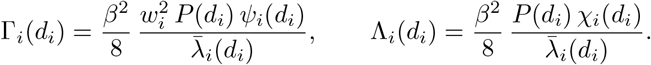

##### Simplified regime and interpretation

In the general formulation, the coupling coefficients *χ*_*i*_(*d*_*i*_) capture how systematic biases in other attribute channels influence the encoding of channel *i*. They enter the Lagrangian as linear terms that can produce small mean shifts in the optimal encoder, effectively coordinating channels under shared information constraints. These terms reflect bias–bias interactions across attributes mediated by the joint distribution *P* (**d**) and the stakes |*d*_*o*_|.

In our experimental setting, however, these effects are expected to be negligible for two reasons. First, one attribute in each type of task for food items (healthfulness in the estimation task, tastiness in the choice task) shows nearly unbiased encoding: the power-law exponent relating internal representation to true difference is close to one, implying *b*_*j*_(*d*_*j*_) ≈ 0 pointwise. This directly eliminates all coupling terms involving that channel, as *χ*_*i*_(*d*_*i*_) depends linearly on *b*_*j*_(*d*_*j*_).

Second, for options with two attributes, *χ*_*i*_(*d*_*i*_) can be treated as negligible or small if the following conditions are met simultaneously: (1) the conditional distribution of *d*_*j*_|*d*_*i*_ is symmetric and centered around its conditional mean 𝔼 (*d*_*j*_|*d*_*i*_); (2) prior *P* (*r*_*j*_) is symmetric around 0; (3) the encoder optimization is sign-symmetric around zero (i.e., the encoding rule treats positive and negative attribute differences equivalently). Condition (2) and (3) are part of the assumptions of our model. For condition (1), we found in both food and social choice datasets, residuals of *d*_*j*_ around the fitted trade-off line *d*_*j*_ = *α* + *βd*_*i*_ had mean magnitudes less than 10% of the attribute scale in all bins of *d*_*i*_ (Supplementary Table S9 and S11, Supplementary Figure S21).

Together, these conditions justify setting the bias–bias coupling term to zero, yielding the simplified encoder:

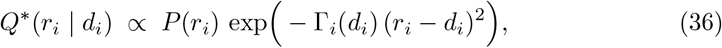

with

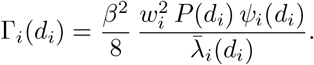

#### Model simulation

We conducted simulations of the goal-adaptive difference encoding model following Cheyette and Piantadosi [77]. Regarding the estimation task model, for each stimulus value, we solved for the Lagrange multiplier *λ*_*d*_*i* that saturates the channel-capacity constraint KL(*Q*, |, *P*) = *C*_*i*_, using gradient descent for 5,000 iterations. The same prior was used for the attribute value *d*_*i*_ and its encoded representation *r*_*i*_.

For the choice task, we assumed two attributes that share a single capacity budget *C*. We therefore fit one shared Lagrange multiplier *λ*(*d*_1_, *d*_2_) for each pair on the full 2-D grid of stimulus values, jointly satisfying KL_1_ +KL_2_ = *C*. To obtain trial-averaged mean responses *E*[*r*_*i*_ | *d*_*i*_], we computed the per-pair mean *E*[*r*_*i*_ | *d*_1_, *d*_2_] under the joint-grid encoder and then marginalized over the off-channel attribute weighted by its prior, *E*[*r*_*i*_ | *d*_*i*_] =∑ *d*−*iE*[*r*_*i*_ | *d*_1_, *d*_2_], *P* (*d*_−*i*_). Both attributes were assigned the same monotonically decaying prior.

### Attribute value difference estimation task

#### Participants

Twenty-seven individuals (52% females and 48% males) participated in this experiment. All participants provided written informed consent in accordance with the procedures of the Institutional Review Board of the Faculty of Business, Economics and Informatics at the University of Zurich and the Ethics Committee of the Canton of Zurich.

#### Task

Participants completed a computer-based food evaluation task. The experiment consisted of two main phases: a rating phase and a difference estimation phase.

In the rating phase, participants viewed images of 50 food items presented individually. Each food item was evaluated separately for its tastiness and healthfulness. The order of rating type (tastiness vs. healthfulness) was randomized across food items. Ratings were made using a continuous scale ranging from –5 (“very bad taste”/”very unhealthy”) to +5 (“very good taste”/”very healthy”), with 0 representing a neutral judgment. Participants selected their responses by clicking with the mouse on the scale. Participants were encouraged to use the full range of the scale across foods. Each food item appeared twice, although participants were not explicitly informed about the repetition. Participants were encouraged to try their best to provide honest ratings to help them achieve better performance in the next phase of the task.

During the attribute value difference estimation phase, participants viewed pairs of food items and indicated which item was tastier, or healthier on separate trials, as well as the perceived magnitude of the difference. The evaluation were made on a continuous scale anchored at the midpoint (no difference). Clicking on the left or right side of the bar indicated which item was judged superior, and the distance from the midpoint indicated the perceived magnitude of the difference. The true difference was derived from participants’ own ratings in the prior phase, and they were informed that more accurate estimates would lead to greater monetary rewards.

### Multi-attribute choice datasets

#### Food-choice dataset Food1

##### Participants

This experiment included 100 participants (25% females and 75% males). Participants were instructed to fast for 2.5 hours before the experiment. Compliance with this instruction was monitored through self-report. All participants provided written informed consent in accordance with the procedures of the Institutional Review Board of the Faculty of Business, Economics and Informatics at the University of Zurich and the Ethics Committee of the Canton of Zurich. In addition to the food they chose, all participants were also given a flat fee to compensate them for their time.

##### Task

The study involved a two-part task, with the first part comprising a rating task in which participants were asked to evaluate the tastiness or healthfulness of each food item in the choice set, with independent blocks for each attribute. Participants provided their rating by indicating their response on a continuous scale with numerical anchors of -5 and 5 at each end.

In the second part, participants were asked to make decisions between two options in a choice task. The choice task consisted of three types of decision conditions, each presenting the food items on the screen in a different manner:

- In the “Ratings” condition, participants were presented with only the healthfulness and tastiness ratings they provided in the previous rating task, without any images of the food items. The tastiness ratings were displayed on the top half of the screen, while the healthfulness ratings were displayed on the bottom half.
- In the “Images” condition, participants were presented with only the images of the two food items, without any accompanying ratings or text.
- In the “Ratings & Images” condition, participants were presented with both the images of the food items and the healthfulness and tastiness ratings they provided in the previous rating task. The tastiness ratings were displayed on the top half of the screen, while the healthfulness ratings were displayed on the bottom half. The images of the two food items were shown in the middle of the two ratings.

All participants completed a total of 180 trials, with 60 trials in each of the three experimental conditions. The trials within each condition were presented in blocks of 8, 12, 18, or 22 trials. The order of the condition blocks was randomized across participants. The same 60 pairs of food options were used in all three conditions. We present our primary analyses of the “Images” condition in the main text because that condition corresponds to the typical food choice presentation format. The results for the other two conditions are similar and are included in the supplementary materials.

#### Food-choice dataset Food2

We used the choice and RT data from the “Low working memory load (LL)” condition in the dataset of [78].

##### Participants

Fifty participants (23 males) aged 18–30 years were recruited for the experiment via online advertisements. All potential participants were screened for exclusion criteria such as contraindications to MRI, a history of mental disorder (including eating disorders), or any neurological illness. Individuals who reported experiencing difficulties with dietary self-control and a general interest in healthy eating were invited to participate in the study. Individuals who adhere to a vegan diet or exclude certain nutrients from their diet for health reasons were not eligible to participate in the study. To ensure consistency in the level of hunger among participants, they were instructed to consume a small meal (e.g., two sandwiches) 2 h prior to the experiment. The authors obtained written consent from each participant. The procedure was approved by the Kozminski University Ethics Committee (No 2, 2019/2020, 30/06/2020).

##### Task

First, the authors collected the subjective healthfulness and tastiness ratings of ≤ 414 food items from each participant using a continuous visual scale, which ranged from 5 (very untasty/unhealthy) through 0 (neutral tastiness/healthfulness) to 5 (very tasty/healthy). The order of the tastiness and healthfulness ratings was counterbalanced across participants. Based on these ratings, they created 90 pairs of food items, which were unique for each participant: 60 challenging pairs (a healthier and less tasty food item and a tastier and less healthy food item) and 30 non-challenging pairs (a healthier and tastier food item and a less healthy and less tasty food item). The number of food products evaluated varied between study participants. The rating process continued until the algorithm was able to create the aforementioned 90 food pairs.

Prior to the choice session, participants agreed to select the healthiest available options whenever possible, while also taking into account their own taste preferences. To ensure the realism of the computerized series of food choices, the participants were informed that one randomly selected food item chosen by them would be provided for consumption after the experiment.

The experiment was conducted in a fMRI scanner. Participants made 90 food choices in two conditions: under low (LL) and high (HL) working memory load. Prior to making each food choice, participants were to memorize a seven-digit number (HL) or a one-digit number (LL) in 3 s. Then they were presented with a pair of food items and had 3.5 s to make a selection. Finally, a number was presented on the screen, and participants were to indicate whether it matched the number they had to memorize. For each correct response, a bonus of 0.01 PLN was added to the participation fee of 120 PLN.

In total, 200 trials were conducted with 100 trials per condition: 120 trials with food choices that required self-control (60 challenging pairs presented twice: in HL and LL conditions); 60 trials with food choices that did not require self-control (30 non-challenging pairs presented twice: in HL and LL conditions); and 20 trials without any food choices (10 trials in each condition with a blank screen instead of a food choice). The order of both load conditions and the types of pairs (challenging and non-challenging) was fully random and unique for each participant.

#### Dictator-game dataset Social1

We used the choice and RT data from the second half of the “time-free” condition in the dataset of [5], same as the implementation as the original paper. These data are openly available at https://doi.org/10.17605/OSF.IO/UKG7B.

##### Participants

In total 102 subjects (56 females) participated in the experiment. Eighteen subjects took part in an initial experiment at The Ohio State University (OSU), followed by 84 subjects at the University of Konstanz. On average, subjects earned 20 dollars at OSU and 16 Euros at Konstanz (including show-up fees). Subjects gave informed written consent before receiving the instructions at OSU, and we obtained informed consent from subjects when they registered for the experiment at Konstanz. OSU’s Human Subjects Internal Review Board approved the experiment.

##### Task

The mini-dictator games under different time conditions had the same properties but minor differences in payoffs. Specifically, the differences between the dictators’ payoffs (DicDiff) were 2, 4, 6, 8, and 10, while the differences between the receivers’ payoffs (ReceDiff) were from 3 to 57, in steps of 6. In every trial, the subject had to decide whether to give up some of their own money in order to increase the other subject’s payoff and reduce the inequality between them. The authors first fixed the parameters for 50 games in the time-free condition (Games 1–50). Then then decreased (increased) all the payoffs by 1 for one half of these games and increased (decreased) all the parameters by 1 for the other half of games to get the 50 games for the time-pressure (-delay) conditions. Finally, they decreased all the parameters by 2 for one half of the games and increased all the parameters by 2 for the other half of games to get the other 50 time-free trials (Games 51–100).

At the beginning of each session, they randomly matched subjects into two-person groups. They randomized the order of the games within the different time conditions for each group.

#### Dictator-game dataset Social2

We used the choice and RT data from the second half of the “time-free” condition in the dataset of [79], same as the implementation as the original paper. These data are openly available at https://pubsonline.informs.org/doi/10.1287/mnsc.2023.4732.

##### Participants

In total, 103 university students (56 females, *mean* = 20.0 years, *sd* = 2.0 years) participated in this experiment from November 20 to December 24, 2021. All participants were right-handed. On average, participants earned 6.4 U.S. dollars (including the show-up fee). The internal review board of Zhejiang University approved the experiment, and all participants provided written informed consent.

##### Task

This is a replication experiment for dataset Social1 [5] with adding additional decision trials, using mouse-tracking instead of button-press responses, and randomizing the assignment of the choice problems to the three time conditions. Specifically, the authors generated another 100 games (two subgroups) using the rules for dataset Social1 [5] besides the original 200 mini-dictator games in [5]. That is, they decreased the payoffs in half of the games in subgroup 1 (with game IDs of 1–50) by two or three and increased the payoffs in the other half of the games by two or three. This ensured that the differences in self and others’ payoffs were identical across the six subgroups. In the experiment, they randomly assigned the six subgroups of games into time-free (four subgroups, 200 games), time-pressure (one subgroup, 50 games), and time-delay (one subgroup, 50 games) conditions at the participant level. Other than creating and randomizing the games across conditions in this manner, they used the same procedures in this experiment as in [5].

### Regression analysis

#### Comparing linear and power-law Bayesian hierarchical models for attribute valuation

We evaluated whether nonlinear power-law models provide a better fit to the data than standard linear regression models to describe the relationship between reported estimations for attribute value differences and true attribute value difference. These models were implemented using the *brms* package in *R*, which fits Bayesian generalized linear mixed-effects models.

The linear model was specified as:

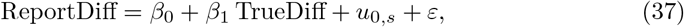

where ReportDiff is the reported estimation of attribute value difference, and TrueDiff is true attribute value difference based on the ratings. Both ReportDiff and TrueDiff are on their original scale, as in the task setting, [-10, 10], unless stated otherwise. The term *u*_0,*s*_ is a subject-specific random intercept and slope across subjects, and *ε* denotes the residual error term.

The nonlinear models incorporated a power parameter *k* and were specified as:

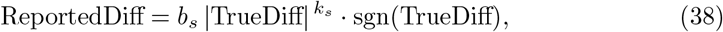

where *b*_*s*_ and *k*_*s*_ are subject-specific parameters defined as

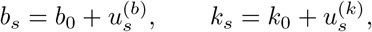

with 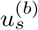 and 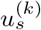 denoting subject-level random effects. To prevent the exponent from changing the sign of the input value, we applied the power-function transformation to the absolute value of TrueDiff and then multiplied i t by the original sign of it.

We used the following priors for the regression models:

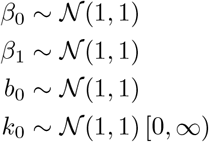

The model comparison procedures follow the section “Model comparison via leave-one-out cross-validation (LOO)” below. The model fitting and comparison were run for each attribute separately. For each model, we used four chains with 2,000 samples per chain after burn-in.

#### Comparing homoscedastic and heteroscedastic linear mixed-effects models for attribute value difference estimation variance

We evaluated whether heteroscedastic linear mixed-effects (LME) models provide a better fit to the estimation task data than standard homoscedastic LME models to describe the variance differences between attributes (Figure 3c and Supplementary Table S8). In the heteroscedastic model, the residual variance was allowed to vary across attribute types (tastiness, healthfulness). The models were implemented using the *nlme* function from the *nlme* package in *R*, utilizing the *varIdent* weight function to model the unequal variances. The standard homoscedastic model was fitted assuming constant residual variance.

The regressors in the models are listed in the equation below:

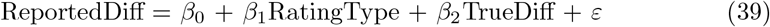

The model included random intercepts and random slopes for subjects. The heteroscedastic version allowed the residual variance to differ by “RatingType”.

#### Analyzing attention allocation with linear mixed-effects model and Beta regression

We evaluated the allocation of visual attention (fixation) toward attribute ratings. To examine the difference in fixation time between tastiness and healthfulness ratings, we fitted a standard linear mixed-effects model using the *lmer* function from the *lmerTest* package in *R*. To analyze the proportion of fixation time spent on tastiness versus healthfulness ratings, we employed a Beta regression model, as the dependent variable was a proportion bounded between 0 and 1. The model was implemented using the *glmmTMB* function from the *glmmTMB* package in *R*, utilizing a Beta family with a logit link function.

For the fixation time difference analysis, the model is described in the equation below:

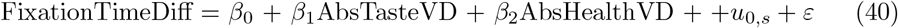

In this model, FixationTimeDiff denotes the difference in fixation time between tastiness and healthfulness ratings. The regressors are the absolute value differences for tastiness (AbsTasteVD) and healthfulness (AbsHealthVD). Unless stated otherwise, both AbsTasteVD and AbsHealthVD are analyzed on the original task scale, [0,10]. FixationTimeDiff is in milliseconds. This model included random intercepts and random slopes for subjects (*u*_0,*s*_).

For the fixation proportion analysis, the regressors are listed in the equation below:

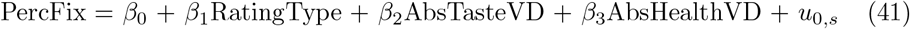

Here, PercFix is the proportion of time spent fixing on a specific rating type. RatingType is a categorical factor distinguishing between healthfulness and tastiness ratings. The model included random intercepts and random slopes for subjects (*u*_0,*s*_).

### Drift-diffusion models (DDM)

Across all DDM variants for all datasets, we implemented hierarchical Bayesian modeling in *JAGS* [80]. We ran three parallel chains, discarding the first 50,000 iterations of each as burn-in and retaining 10,000 thinned posterior samples per chain.

For the Food1 dataset, we used a three-level hierarchical structure to improve stability given the relatively small number of trials per condition. Specifically, parameters were estimated at the population level across all subjects, with subject-level parameters modeled as draws from the population distribution. Within each subject, condition-specific parameters were modeled as draws from the corresponding subject-level distributions, yielding separate parameter estimates for each condition while enabling partial pooling across conditions and subjects.

For the other datasets, we used a simpler hierarchical structure with a single hierarchy across subjects (i.e., population-level priors with subject-level parameters), without an additional condition level.

#### Baseline DDM with weighted additive decision rule (WAD)

In the baseline DDM with a weighted additive decision rule (WAD), the drift rate (*v*) is modeled as a linear combination of the value differences for two attributes—*d*_1_ and *d*_2_—weighted respectively with *w*_1_ and *w*_2_. For the dictator-game datasets, we additionally included a constant term (*c*) in the drift rate equation, as prior work has shown this improves model fit [5]. The drift rate is defined by the following equation:

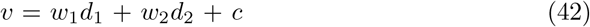

We also incorporated a relative starting point bias (*x*_0_) and non-decision time *NDT* in the baseline model. During model estimation, the decision boundary was held constant, while the noise parameter (*σ*) was treated as a free parameter.

We used the following priors for the population-level parameters:

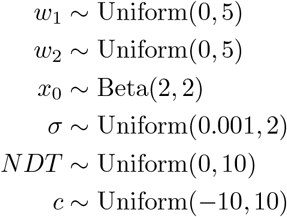

For the DDMs applied to the food dataset, we additionally incorporated loss aversion in the tastiness attribute by linearly amplifying negative tastiness values. This modeling choice was motivated by clear discrepancies in choice behavior for positive versus negative tastiness values (Figure S7a), as well as by the results of the model comparison analyses (Tables S1, S2, and S3). In contrast, no such discrepancy was observed for the positive and negative healthfulness values (Figure S7b), therefore loss aversion was not modeled for the healthfulness attribute. Loss aversion in tastiness was implemented following the approach described in [81]:

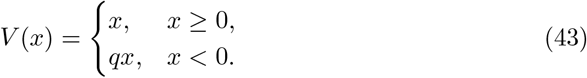

In this formulation, *x* represents the original tastiness rating, and *q* captures the degree of loss aversion. A value of *q* = 1 corresponds to no loss aversion, while values greater than 1 indicate amplification to n egative tastiness r atings. D uring m odel fitting, *q* was constrained to be greater than or equal to 1. We used the following priors for the population-level loss aversion parameter, with the other parameters following the priors of the baseline DDM:

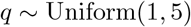

For the food-choice models, the input variables were kept on the original rating scales specified in the task description and were then subjected to either a loss-aversion transformation or a power-function transformation, depending on the model. For the social-choice models, the input variables were rescaled by dividing by 10 relative to the original payoff scales described in the task.

#### DDM with power-function transformation of attribute value differences

With power-function transformation of attribute value differences, the corresponding drift rate equation is as follows:

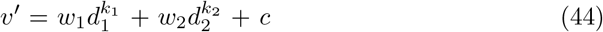

Similar to the practice in regression models, to prevent the exponent from changing the sign of the input value, we applied the power-function transformation to the absolute value and then multiplied it by the original sign. The baseline DDM is equivalent to the case where both *k*_1_ and *k*_2_ are equal to 1.

We used the following priors for the population-level exponent parameters, with the other parameters following the priors of the baseline DDM:

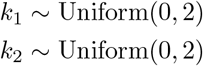

For the model incorporating both loss aversion on tastiness and power-function transformation on attribute values, the tastiness rating is firstly transformed with loss aversion, then goes through power function transformation on the value difference.

#### DDM model simulation

We generated simulated choices by parameterizing the DDM using the median of the fitted posterior probability chains of the parameters. The simulated choices were based on these parameters and the attribute values participants faced during every trial. Because there is randomness due to the noise parameter in the DDM process, for plotting the simulated data, we generated 10 simulations and took the average of them.

### Bayesian hierarchical model comparison via leave-one-out cross-validation (LOO)

We implemented the model comparison using the “loo” package in *R* [28], which is based on ELPD (Expected Log Pointwise Predictive Density) representing the theoretical ELPD for a new dataset. ELPD can be estimated using methods such as cross-validation. The term “elpd_loo” in the model comparison refers to the Bayesian leave-one-out estimate of the expected log pointwise predictive density and is calculated as the sum of N individual pointwise log predictive densities.

Subsequently, this method computes the values of “elpd diff” and “se diff” by performing pairwise comparisons between each model and the model with the largest ELPD (the model in the first row in the model comparison table).

## Acknowledgments

We thank Gaia Lombardi for her efforts in designing and collecting data for the Food1 study, and the authors of the original papers for sharing data from Food2, Social1, and Social2. We also thank Cendri Hutcherson, Nikki Sullivan and Kenway Louie for their comments on this study, and Samuel J. Cheyette for code sharing and suggestions on model fitting.

SDB, SB and DL gratefully acknowledge the support of Marlene-Porsche Foundation scholarships for their PhD studies. SDB was additionally funded by UZH Candoc Grant for her PhD studies. This work was funded in part by the European Union 7th Framework Programme grant number 607310 (NUDGE-IT; site PI: TAH). The acquisition of the Food2 dataset was supported by a grant from the National Science Centre, Poland ‘Cognitive resources and self-control of impulses: a neuroeconomic approach’ (No. 2017/26/D/HS6/01157; PI: L-ukasz Tanajewski).

## Supplementary Figures

**Fig. S1.**
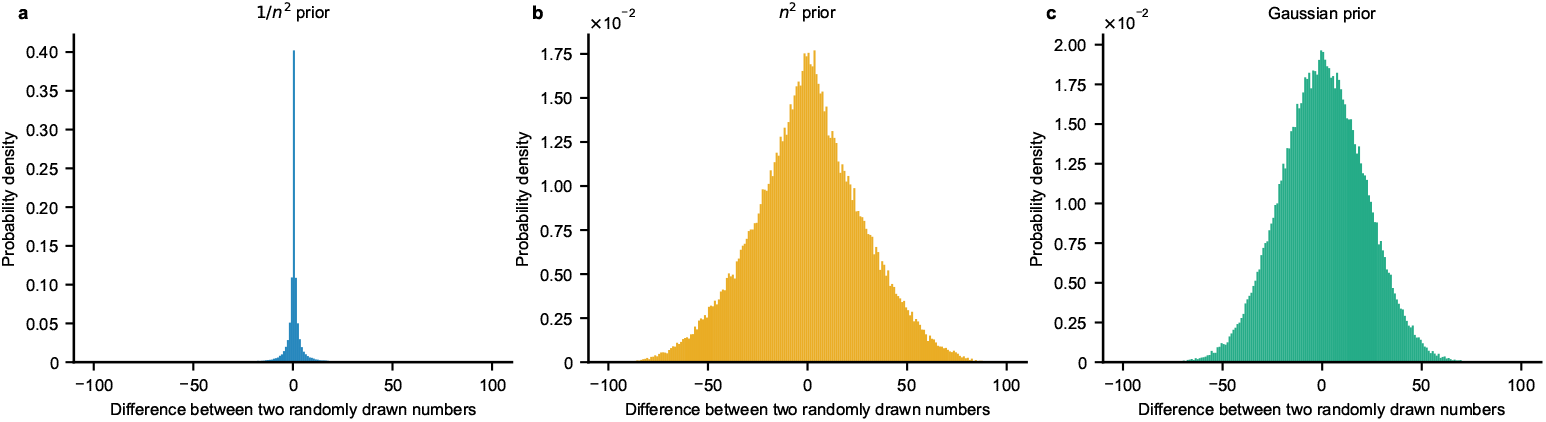
Priors of different shape over individual values can all imply decaying difference distributions. In panels a-c, the histograms show the distribution of pairwise differences obtained by randomly drawing two values from priors of different shape over individual values and subtracting them. The shape of the individual value distributions for each panel is given in the panel titles. In all cases, the resulting difference distributions are peaked near zero and exhibit decaying tails.

**Fig. S2.**
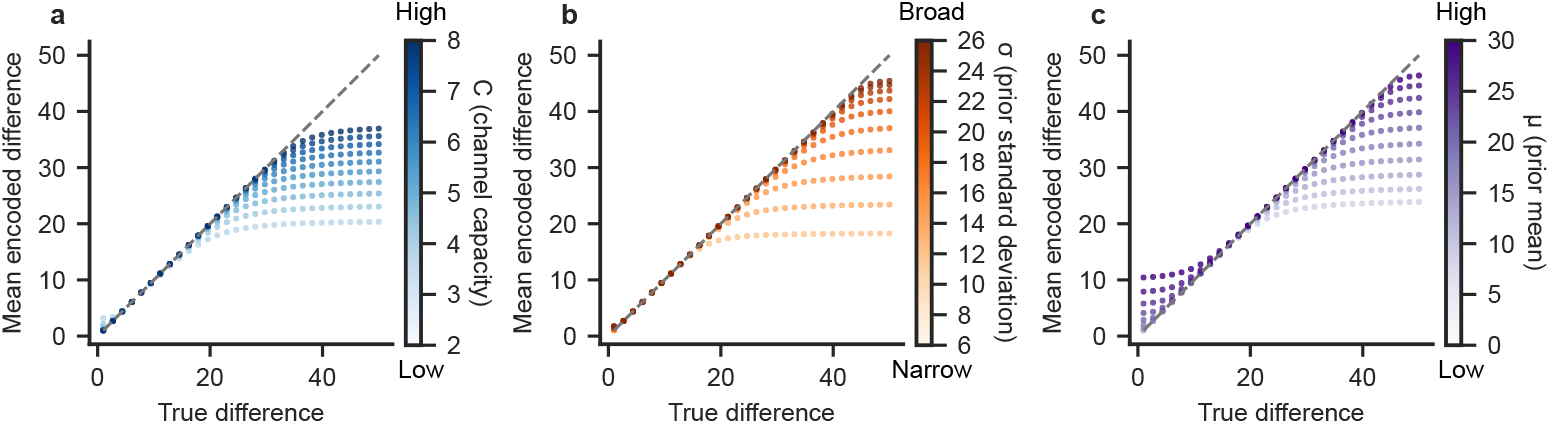
Predictions for the estimation task using a truncated Gaussian prior. This figure generalizes the pattern shown in main text Figure 1 from a power-law to a Gaussian prior shape. We vary the channel capacity parameter *C*, while keeping the prior fixed with mean *µ* = 5 and standard deviation *σ* = 10. Higher *C* corresponds to higher capacity and produces responses closer to the identity line (exponent closer to 1 in the fitted power function). We used 10 evenly spaced values of *C* ∈ [2, 8] for the simulation. (b) We fix *C* = 4 and vary the standard deviation *σ*. Smaller *σ* yields narrower priors and therefore stronger compression. We used 10 evenly spaced values of *σ* ∈ [6, 26] for the simulation. (c) We fix *C* = 4 and vary the mean *µ*. We used 10 evenly spaced values of *µ* ∈ [0, 30] for the simulation.

**Fig. S3.**
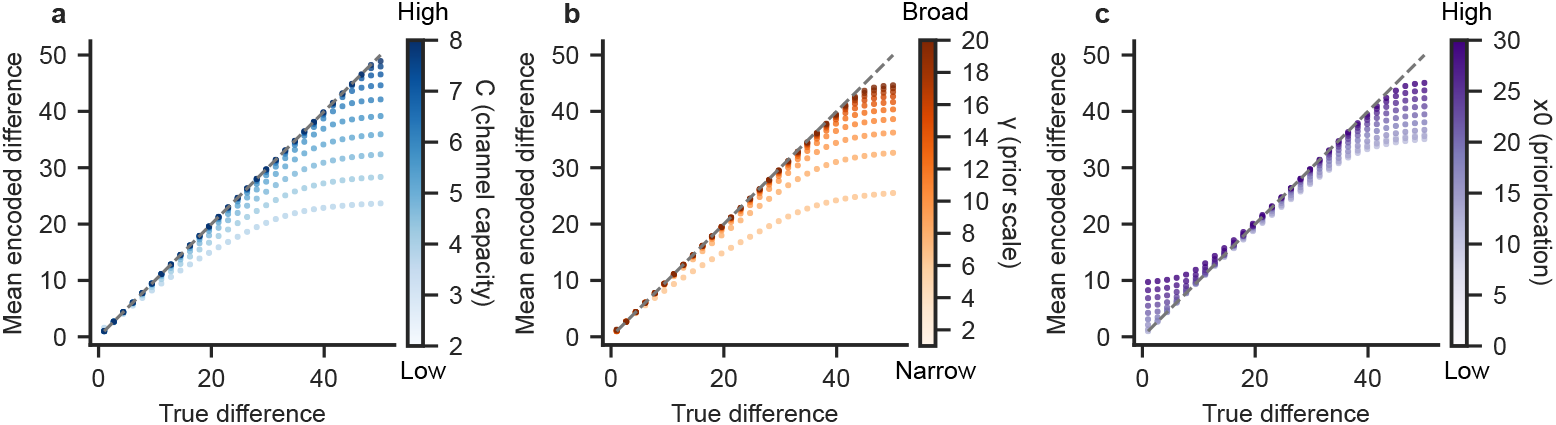
Predictions for the estimation task using a truncated Cauchy prior. This figure generalizes the patterns shown in main text Figure 1 and Figure S2 to a Cauchy prior shape. (a) We vary the channel capacity parameter *C*, while keeping the prior fixed with location *x*0 = 0 and scale *γ* = 5. Higher *C* corresponds to higher capacity and produces responses closer to the identity line (exponent closer to 1 in the fitted power function). We used 10 evenly spaced values of *C* ∈ [2, 8] for the simulation. (b) We fix *C* = 4 and vary the scale *γ*. We used 10 evenly spaced values of *σ* ∈ [1, 20] for the simulation. (c) We fix *C* = 4 and vary the location *x*0. We used 10 evenly spaced values of *x*0 ∈ [0, 30] for the simulation.

**Fig. S4.**
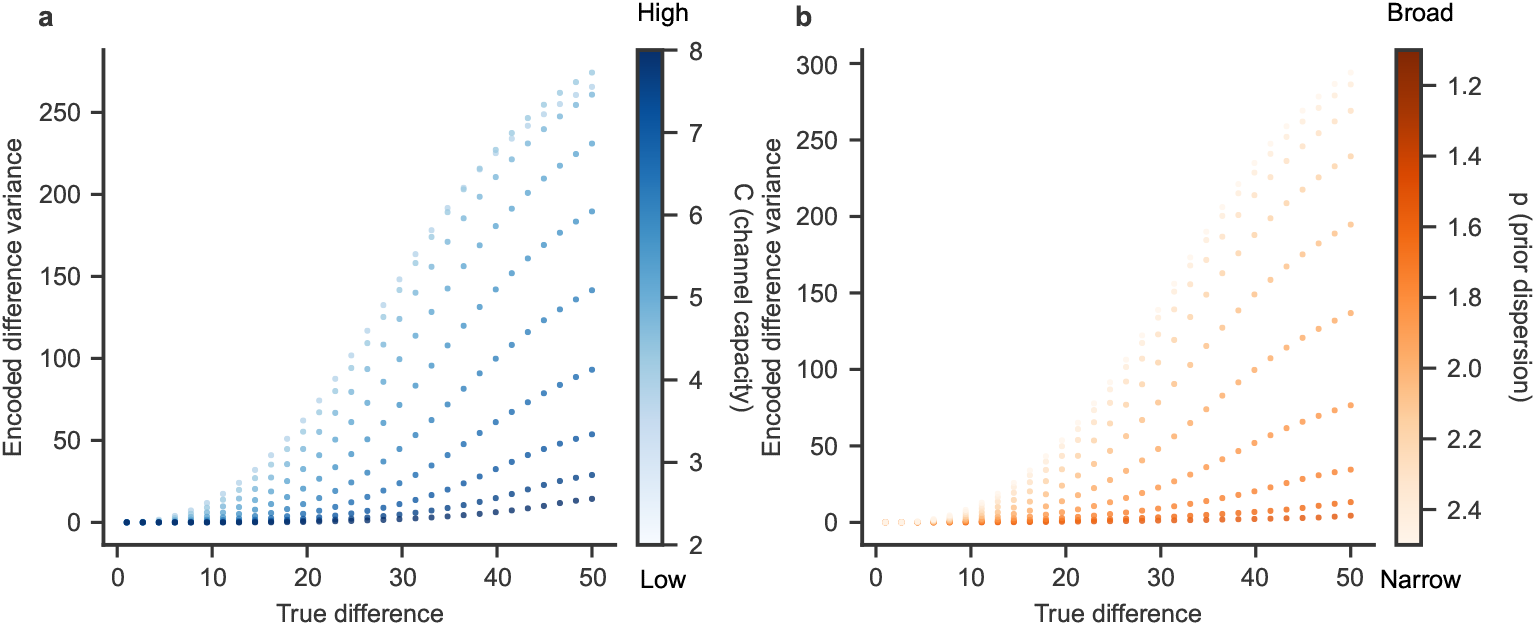
Channel capacity and prior dispersion jointly determine posterior variance in the estimation task. Simulations from the goal-adaptive difference encoding model show the posterior variance of the encoded difference responses, Var[*r*_*i*_ | *d*_*i*_], as a function of the true attribute value difference. **(a)** Channel capacity *C* was varied (*C* ∈ [2, 8]) while holding the prior fixed to a power-law form with exponent *p* = 2.0. For sufficiently high capacities, posterior variance decreases monotonically with capacity. **(b)** Channel capacity was fixed (*C* = 4) and the dispersion of the prior was varied by changing the exponent parameter *p* in *P* (*d*) ∝ (1 + |*d*|^*p*^)^*−*1^ (*p* ∈ [1.1, 2.5]). Broader priors (smaller *p*) produce higher posterior variance, while narrower priors reduce uncertainty.

**Fig. S5.**
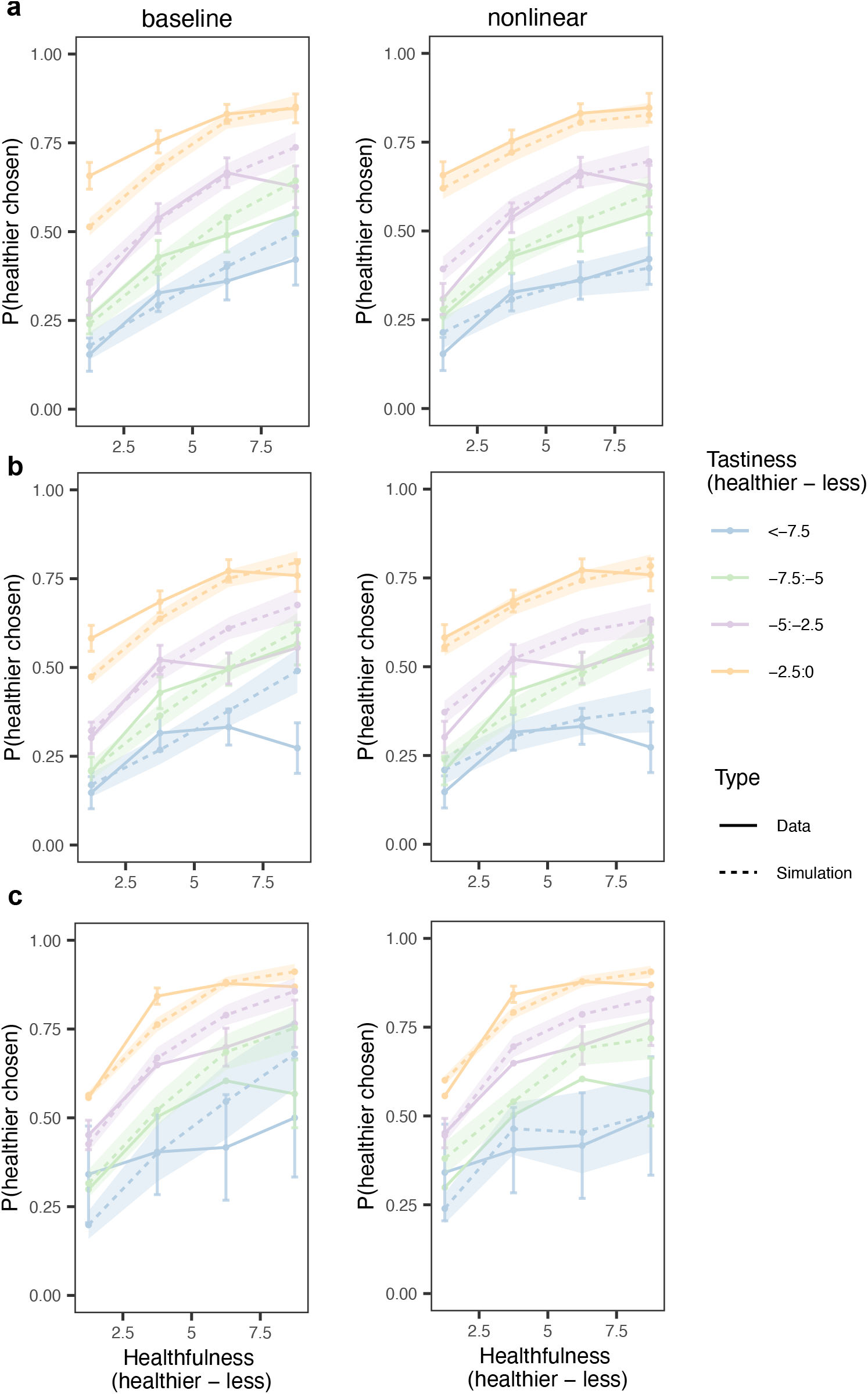
Nonlinear drift rates better capture multi-attribute choice in food decisions. This figure supplements Figure 4 by showing results for additional conditions in the Food1 and Food2 datasets. In all three rows the plots show the probability of choosing the healthier option (y-axis) as a function of the healthfulness difference (healthier option minus less healthier option) between options (x-axis), shown separately for bins of tastiness difference (healthier option minus less healthier option; color coded). Solid lines denote empirical data, and dashed lines denote model simulations. Left panels show simulations from the baseline linear model and right panels show simulations from the nonlinear model. (a) This row shows the results from the Ratings condition from the Food1 dataset in which participants made choices over pairs of foods represented only by the participant’s own previous ratings of taste and healthfulness for each option rather than pictures of the options. This row shows the results from the Ratings & Images condition in the Food1 dataset in which participants made choices over pairs of foods represented by the participant’s own previous ratings and pictures of the two options. (c) This row shows the results of an independent food choice dataset (Food2). The Food2 data come from a traditional presentation format showing pictures of the two food options.

**Fig. S6.**
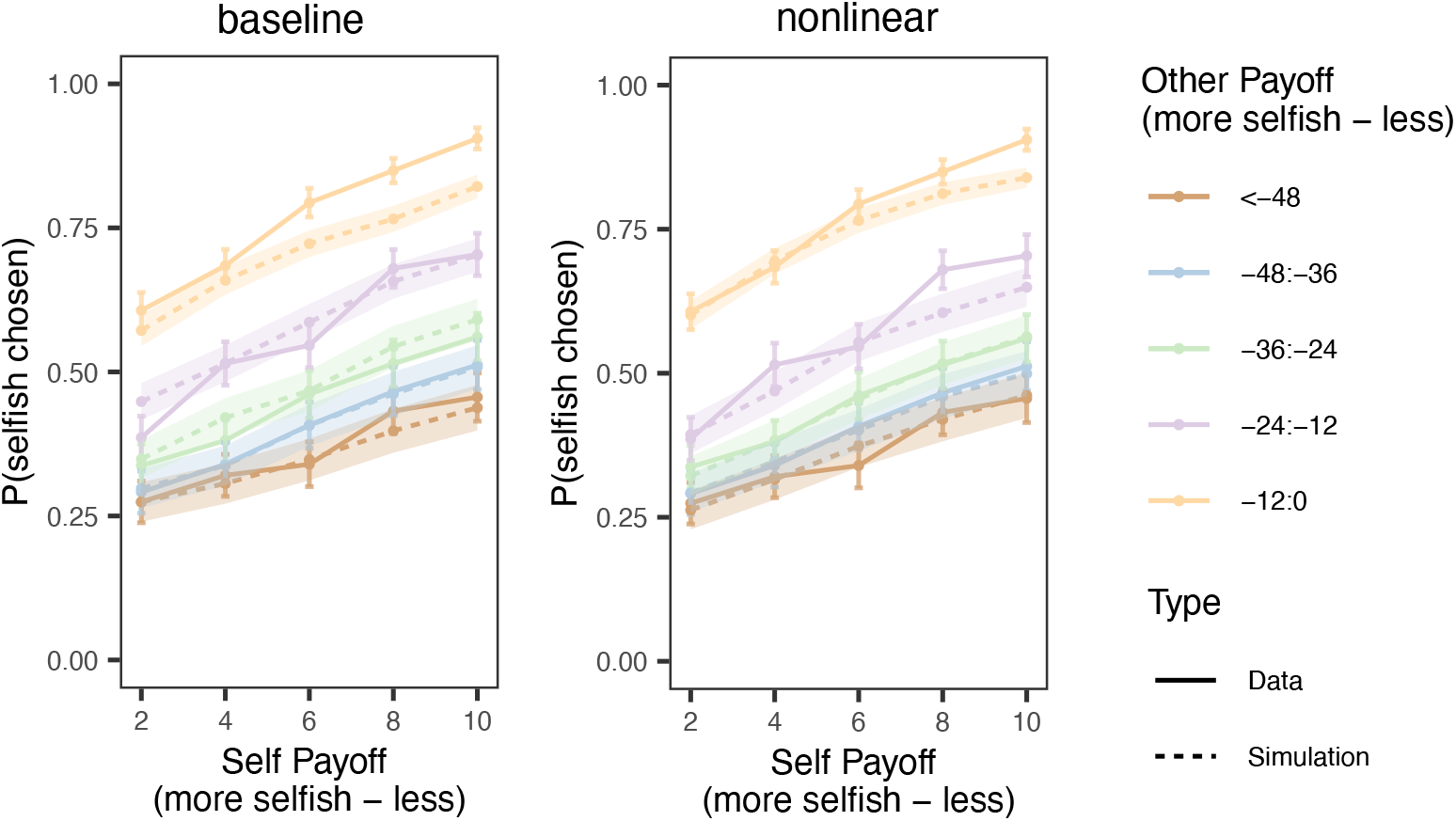
Nonlinear drift rates better capture multi-attribute choice in social decisions in the Time-free condition of the Social2 dataset. This figure supplements Figure 4 by showing the corresponding results for Social2 dataset. The y-axis shows the probability of choosing the more selfish option as a function of self-payoff differences (more selfish option minus less selfish option; x-axis), plotted across bins of other-payoff differences (more selfish option minus less selfish option; color coded).

**Fig. S7.**
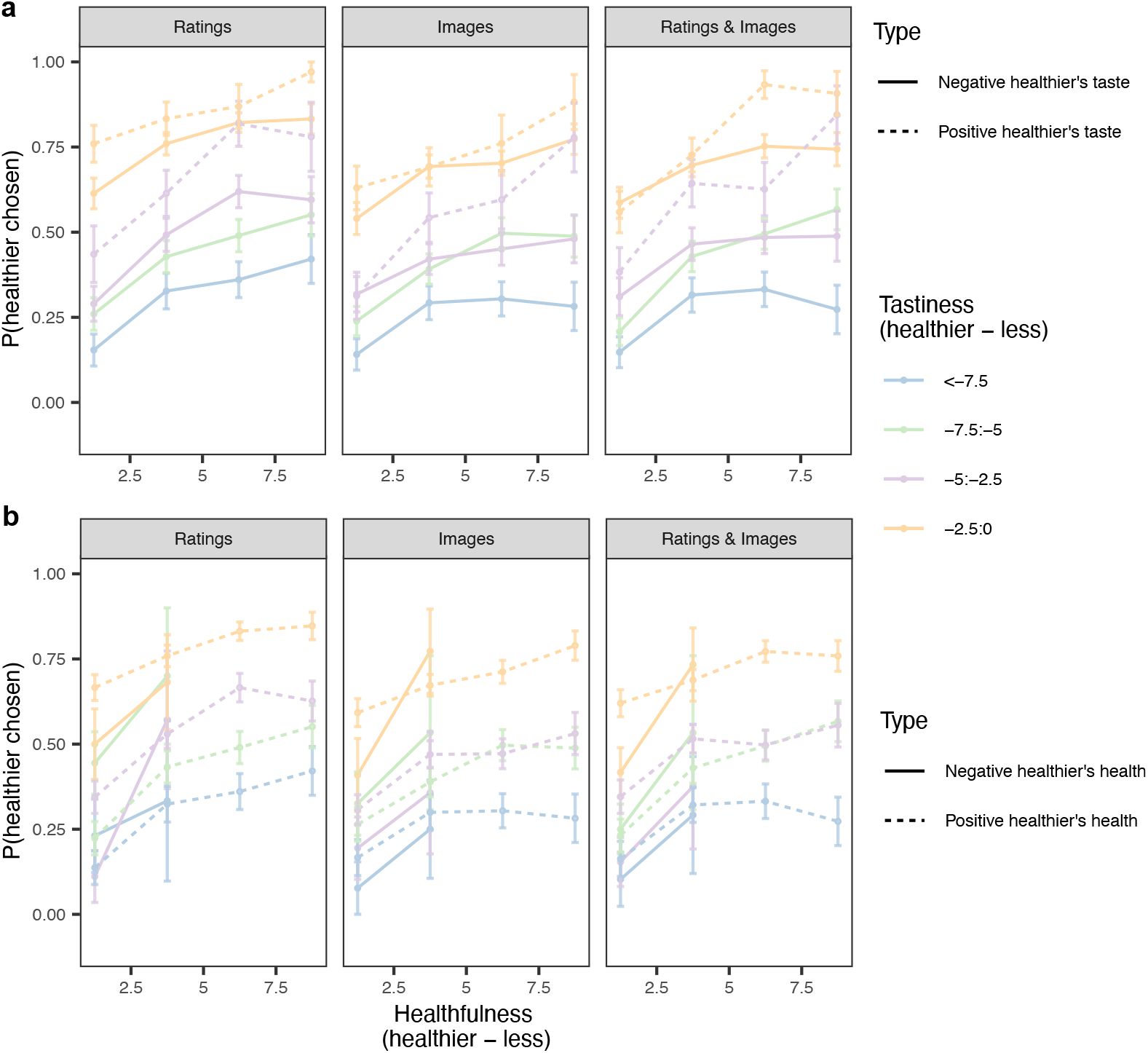
Evidence for loss aversion in tastiness, but not healthfulness, in Food1 dataset. Probability of choosing the healthier option as a function of the healthfulness difference (healthier option minus less healthy option), plotted separately by bins of tastiness difference (healthier option minus less healthy option) and by the sign of the healthier option’s tastiness rating (solid: negative; dashed: positive). Across different conditions, probability of choosing the healthier option is systematically higher for positive tastiness values than that for negative tastiness values. (b) The same analysis for the healthier option’s healthfulness rating shows no systematic asymmetry between positive and negative values, motivating a loss-aversion term for tastiness but not for healthfulness.

**Fig. S8.**
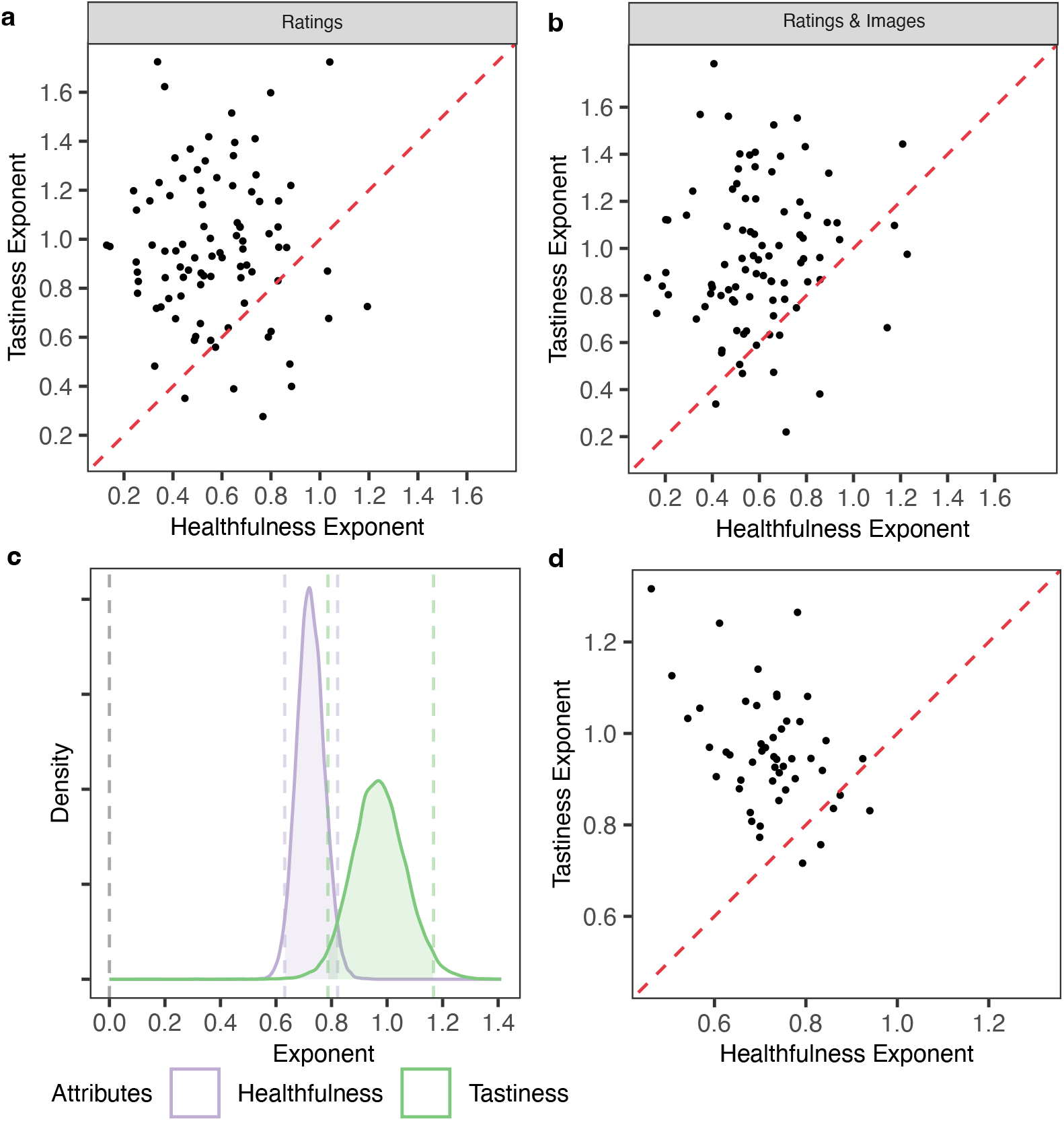
Attribute-specific nonlinearity across food datasets. This figure supplements Figure 5 by showing results for additional conditions in the Food1 and Food2 datasets. (a) Subject-level median exponents for tastiness versus healthfulness in the Ratings condition of Food1 dataset.(b) Subject-level median exponents for tastiness versus healthfulness in the Ratings&Images condition of Food1 dataset. (c) Population-level posterior distributions of exponents in the Food2 dataset. (d) Subject-level median exponents for tastiness versus healthfulness in Food2 dataset.

**Fig. S9.**
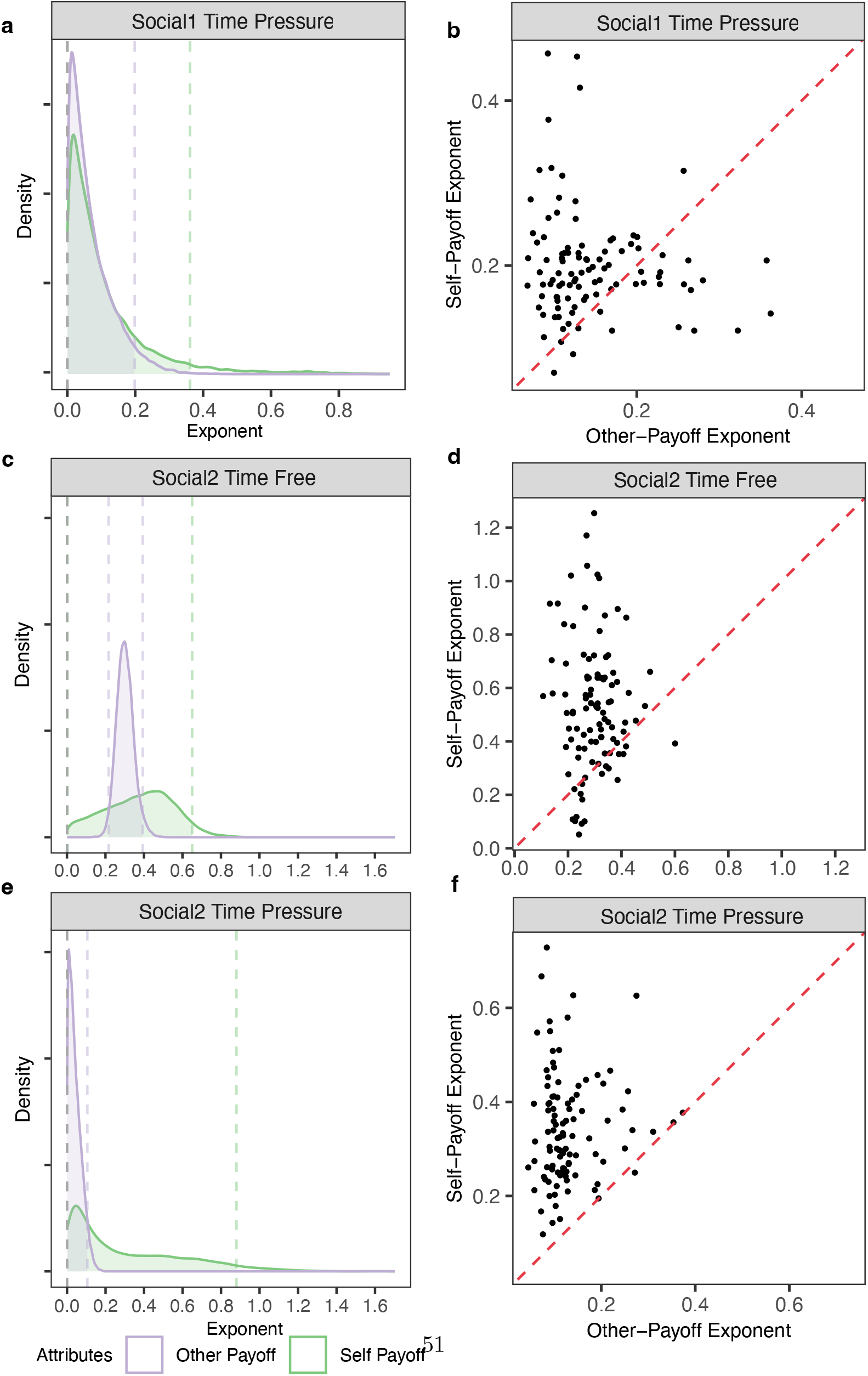
Attribute-specific nonlinearity across social datasets. This figure supplements Figure 5c & d, which show the results of the Time-free condition in the Social1 dataset by showing results for additional conditions in the Social1 and Social2 datasets. (a) Population-level posterior distributions of exponents in Time-pressure condition of Social1 dataset.(b) Subject-level median exponents in Time-pressure condition of Social1 dataset. (c) Population-level posterior distributions of exponents in Time-free condition of Social2 dataset.(d) Subject-level median exponents in Time-free condition of Social2 dataset. (e) Population-level posterior distributions of exponents in Time-pressure condition of Social2 dataset.(f) Subject-level median exponents in Time-pressure condition of Social2 dataset.

## Supplementary Information

### Simulations for effects of stake functions

Recall that the optimal encoding function in a choice task is

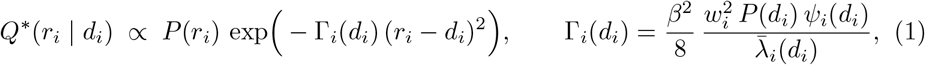

where *ψ*_*i*_(*d*_*i*_) the expected stakes given *d*_*i*_.

In the main text, we used a constant stakes function (*ψ*_*i*_(*d*_*i*_) = 1). Here we show that the main qualitative effects of *C* and *w*_*i*_ hold under more general assumptions about the shape of *ψ*_*i*_(*d*_*i*_) (Figure S10 and Figure S11). Together, these supplementary simulations confirm that the qualitative behavior of the model is robust. While the precise curvature of the encoding function depends on *ψ*(*d*), the key theoretical insight remains unchanged.

**Fig. S10.**
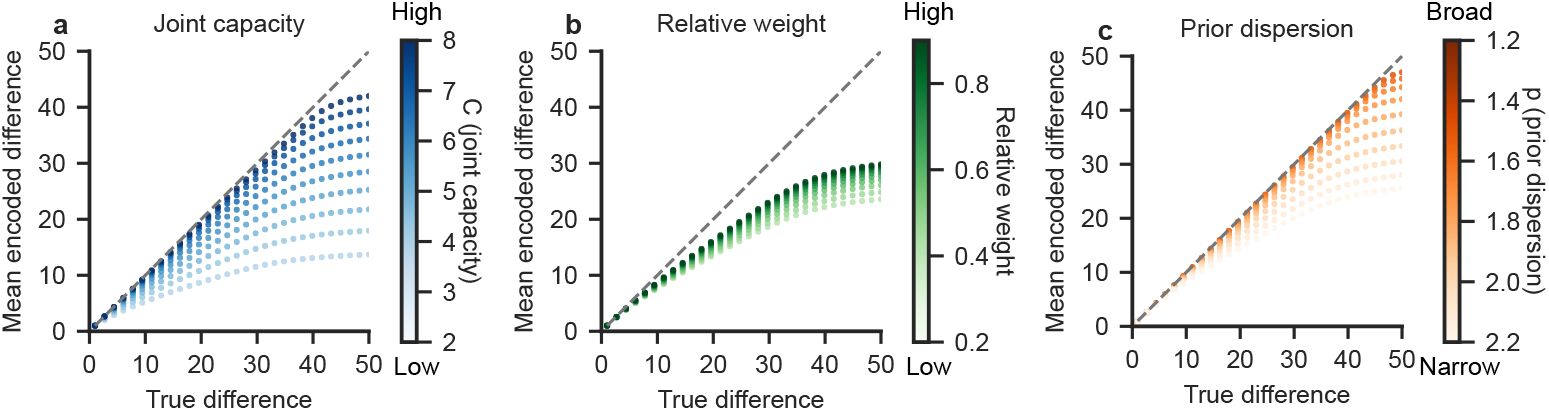
Effect of joint channel capacity and subjective weight in the choice task with stakes increasing with value difference. Simulations show the mean encoded difference responses as a function of the true difference when the expected stakes increase with magnitude. The stakes function was specified as linearly increasing, *ψ*(*d*) = *a* + *b*|*d*| with *a* = 1.0 and *b* = 0.2, such that errors for larger attribute differences are more consequential. **(a)** Channel capacity *C* was varied (*C* ∈ [2, 8]) while holding the subjective weight fixed (*w* = 1.0). Lower capacity leads to stronger compression, while increasing stakes sharpen encoding for larger differences. **(b)** Subjective weight *w* was varied (*w* ∈ [0.2, 0.9]) with channel capacity fixed (*C* = 5). Higher weights reduce compression by increasing encoding precision, particularly at larger magnitudes. **(c)** Prior dispersion *p* was varied (*p* ∈ [1.2, 2.2]) with channel capacity fixed (*C* = 5) and subjective weight *w* = 1.0. For all the simulations, the inverse temperature was fixed at *β* = 1.0, the prior over attribute differences followed an inverse-square form, and the dashed line indicates the identity mapping. These results show that the qualitative effects of channel capacity and subjective weighting on nonlinear encoding are robust to moderate increases in stakes.

**Fig. S11.**
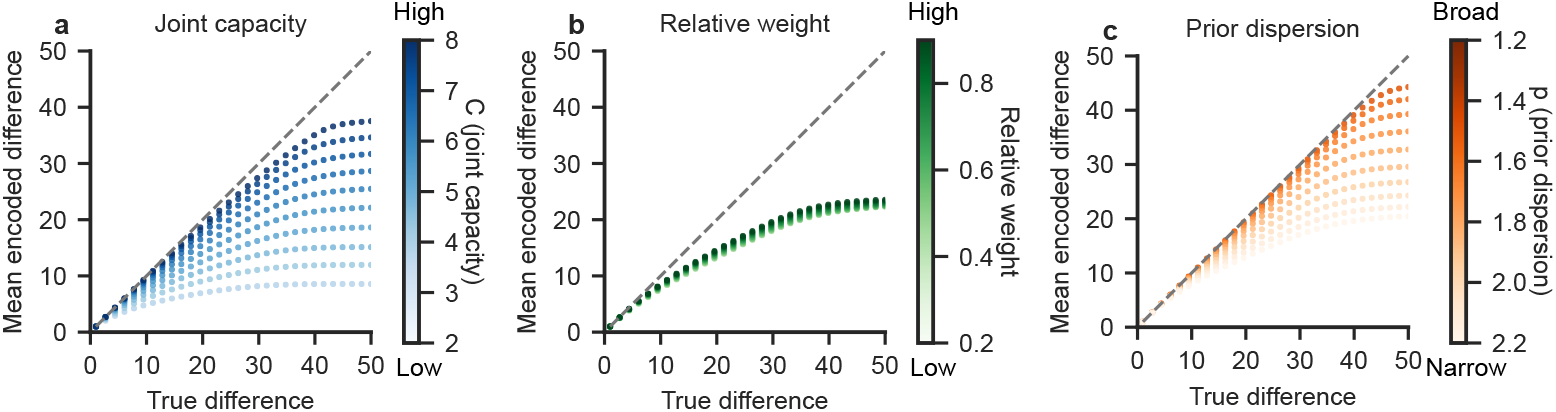
Effect of joint channel capacity and subjective weight in the choice task with stakes decreasing with value difference. Simulations show the mean encoded attribute value difference as a function of the true difference when the expected stakes decrease with magnitude. The stakes function was specified as linearly decreasing, *ψ*(*d*) = *a* − *b*|*d*| with *a* = 3.0 and *b* = 0.2 (truncated at a small positive value to ensure *ψ*(*d*) *>* 0 over the simulated range), such that errors for larger attribute differences are less consequential. **(a)** Channel capacity *C* was varied (*C* ∈ [2, 8]) while holding the subjective weight fixed (*w* = 1.0). Lower capacity leads to stronger compression, with reduced encoding precision at larger differences under decreasing stakes. **(b)** Subjective weight *w* was varied (*w* ∈ [0.2, 0.9]) with channel capacity fixed (*C* = 5). Higher weights reduce compression by increasing encoding precision, though the decreasing-stakes term attenuates precision for larger magnitudes. **(c)** Prior dispersion *p* was varied (*p* ∈ [1.2, 2.2]) with channel capacity fixed (*C* = 5) and subjective weight *w* = 1.0. For all the simulations, the inverse temperature was fixed at *β* = 1.0, the prior over attribute differences followed an inverse-square form, and the dashed line indicates the identity mapping. These results show that the qualitative effects of channel capacity and subjective weighting on nonlinear encoding persist when stakes decrease with value difference.

### Choice model comparisons

**Table S1.** Model comparison including the model with loss aversion for the “Images” condition in Food1 dataset.

|  | elpd_diff | se_diff |
| --- | --- | --- |
| nonlinear model | 0 | 0 |
| nonlinear model without loss aversion | -62.1 | 8.9 |
| linear model | -76.3 | 14.2 |
| linear model without loss aversion | -120.1 | 16.4 |

**Table S2.** Model comparison for “Ratings” condition in Food1 dataset.

|  | elpd_diff | se_diff |
| --- | --- | --- |
| nonlinear model | 0 | 0 |
| nonlinear model without loss aversion | -82.9 | 9.5 |
| linear model | -148.3 | 19.1 |
| linear model without loss aversion | -212.8 | 21.1 |

**Table S3.**
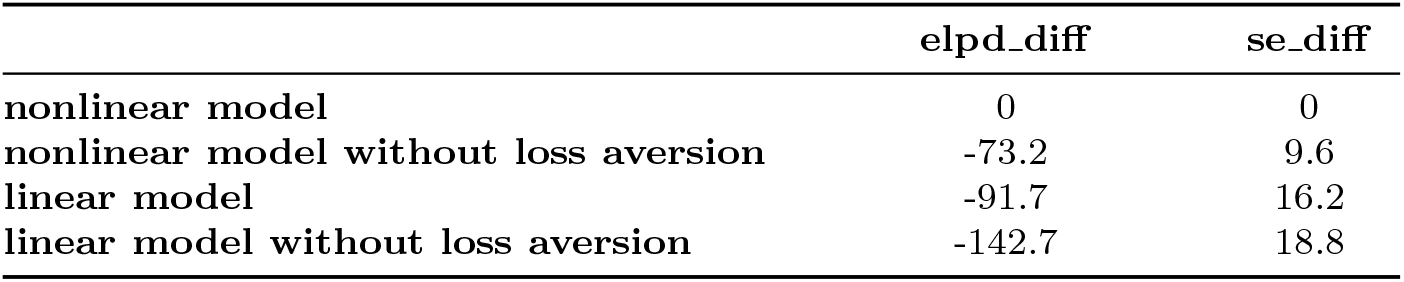
Model comparison for “Ratings & Images” condition in Food1 dataset.

**Table S4.** Model comparison for Food2 dataset.

|  | elpd_diff | se_diff |
| --- | --- | --- |
| nonlinear model without loss aversion | 0.0 | 0.0 |
| nonlinear model | -3.7 | 2.6 |
| linear model | -27.0 | 8.4 |
| linear model without loss aversion | -30.1 | 8.5 |

**Table S5.** Model comparison for “Time-pressure” condition in Social1 dataset.

|  | elpd_diff | se_diff |
| --- | --- | --- |
| nonlinear model | 0.0 | 0.0 |
| linear model | -23.2 | 6.5 |

**Table S6.** Model comparison for the “Time-free” condition in Social2 dataset.

|  | elpd_diff | se_diff |
| --- | --- | --- |
| **nonlinear model** | 0.0 | 0.0 |
| **linear model** | -111.8 | 14.1 |

**Table S7.** Model comparison for the “Time-pressure” condition in Social2 dataset.

|  | elpd_diff | se_diff |
| --- | --- | --- |
| **nonlinear model** | 0.0 | 0.0 |
| **linear model** | -25.7 | 6.4 |

### Testing nonlinearity in multi-attribute choices using segmented regression

In addition to the choice model comparison, we tested the nonlinearity in multi-attribute choices using segmented logistic regression. Using this method does not require a functional form for the nonlinearity. Specifically, we applied both a standard logistic regression model and a segmented logistic regression model to determine whether the latter provides a better description of the empirical or baseline model-simulated data (i.e., simulations from a DDM based on linear attribute value differences).

In the segmented logistic regression model, all the regressors were modeled as having segmented-linear relationships with the response variable, with a single breakpoint estimated for each. The rationale for selecting one breakpoint per regressor was to test whether a minimal degree of nonlinearity could improve the model fit. The model was implemented using the *segmented* function from the *segmented* package in *R*. The standard logistic regression model was fitted using the *glm* function in *R*. After both the standard and segmented logistic regression models were fitted, we conducted model comparison between them using the Bayesian Information Criterion (BIC).

For the food-choice datasets, the regressors in both types of regression models are listed in the equation below:

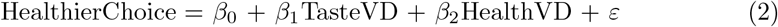

Here, “HealthierChoice” is a binary variable indicating whether the participant selected the healthier option. “TasteVD” and “HealthVD” represent the differences in tastiness and healthfulness ratings, respectively, between the healthier and less healthy options. Both variables were retained on the original rating scales specified in the task description (10 to 10 for TasteVD and 0 to 10 for HealthVD).

For the dictator-game datasets, the regressors in both types of regression models are listed in the equation below:

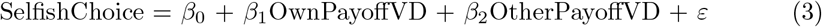

In this case, “SelfishChoice” indicates whether the more selfish option was chosen. “OwnPayoffVD” and “OtherPayoffVD” denote the differences in self’s and other’s payoffs between the more and less selfish options. Both variables were rescaled by dividing by 10 relative to the original payoff scales described in the task.

The segmented logistic regression model provided a better fit to the empirical data, but did not outperform the standard logistic model when applied to the baseline model-simulated data (Figure S12). This result shows that the empirical data has nonlinearity across the attribute value differences, while the baseline choice model specification generates linear choice patterns across the attribute value differences.

**Fig. S12.**
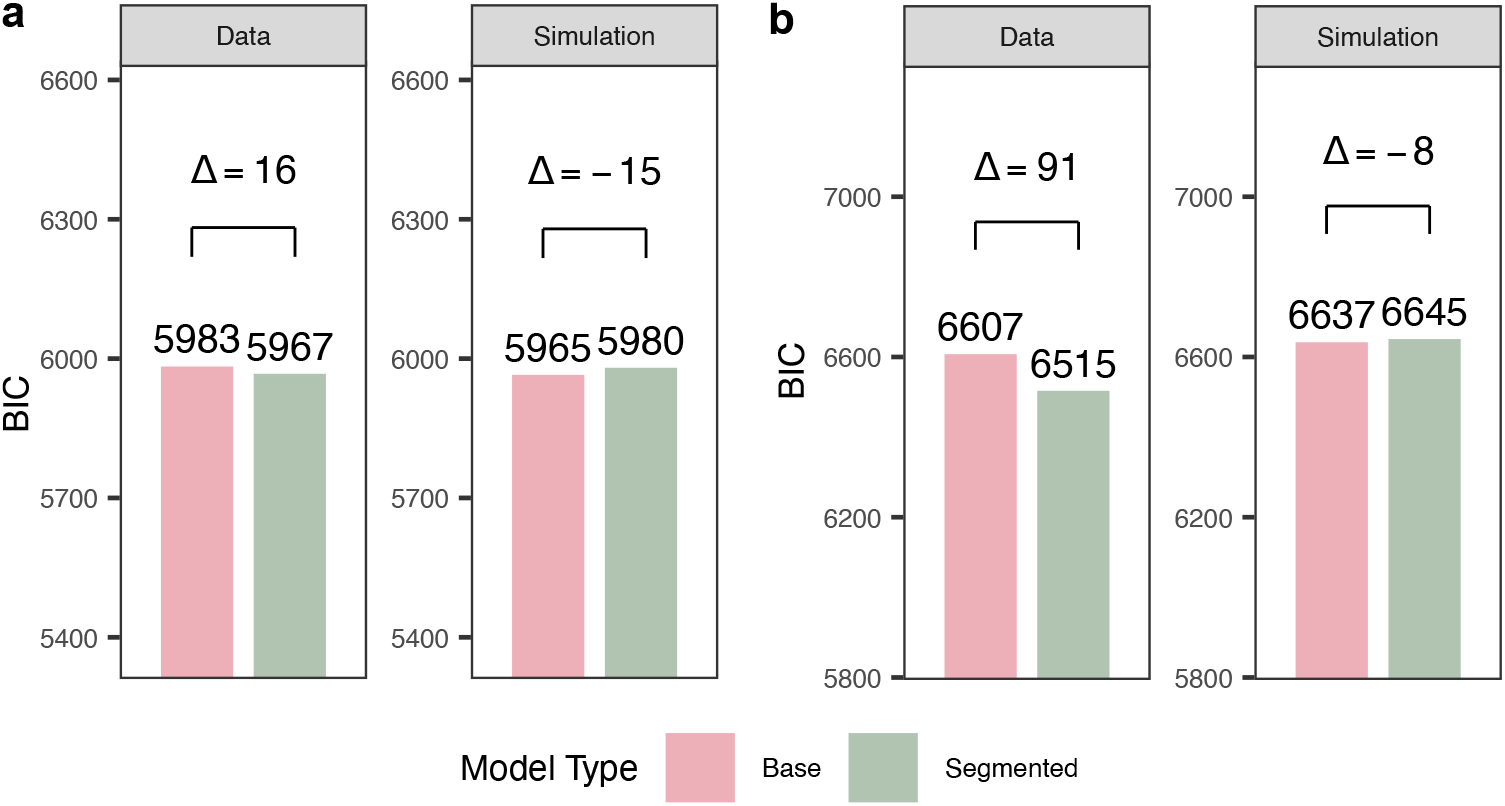
Segmented logistic regression reveals nonlinearity in empirical choices but not in baseline choice model simulations. Comparison of standard versus segmented logistic regression models using Bayesian Information Criterion (BIC). Lower BIC indicates better model fit. (a) Food-choice dataset (Images condition of Food1 dataset). For empirical data (left), the segmented model yields substantially lower BIC than the standard logistic regression (ΔBIC = 16 in favour of the segmented model). In contrast, for data simulated from the baseline choice model (right), the standard logistic model fits as well or better than the segmented model (ΔBIC = 15). (b) Dictator-game dataset (Time-free condition of Social1 dataset). The same pattern is observed: the segmented model improves fit to empirical data (ΔBIC = 91) but not to baseline choice model simulations (ΔBIC = 8). Together, these results demonstrate that real choices exhibit nonlinear dependence on attribute differences, whereas the standard linear model does not.

### Parameter recovery for nonlinear DDM

This section shows the correspondence between the parameters’ posterior distributions after fitting the various empirical datasets (“fitted”, red) and the posterior distributions for parameters fit to data simulated from the model using the parameter values from the initial fits to the empirical data (“Refitted”, blue). High levels of overlap in the red and blue posterior distributions indicate successful parameter recovery.

**Fig. S13.**
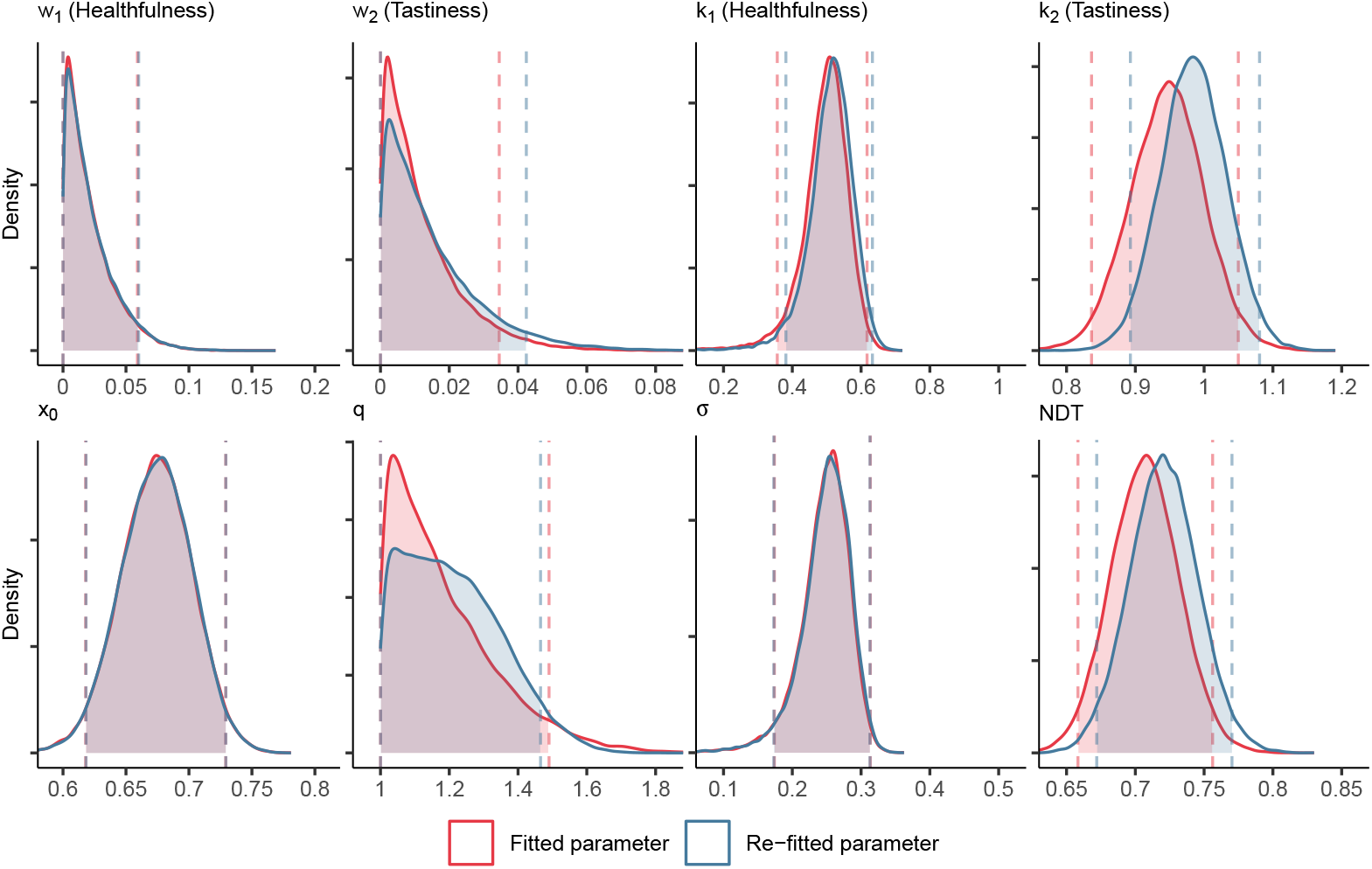
Parameter recovery for the nonlinear DDM with loss aversion in the Food1 dataset. Here and in all subsequent parameter recovery plots, the displayed distributions correspond to the posterior distributions of the population-level parameters.

**Fig. S14.**
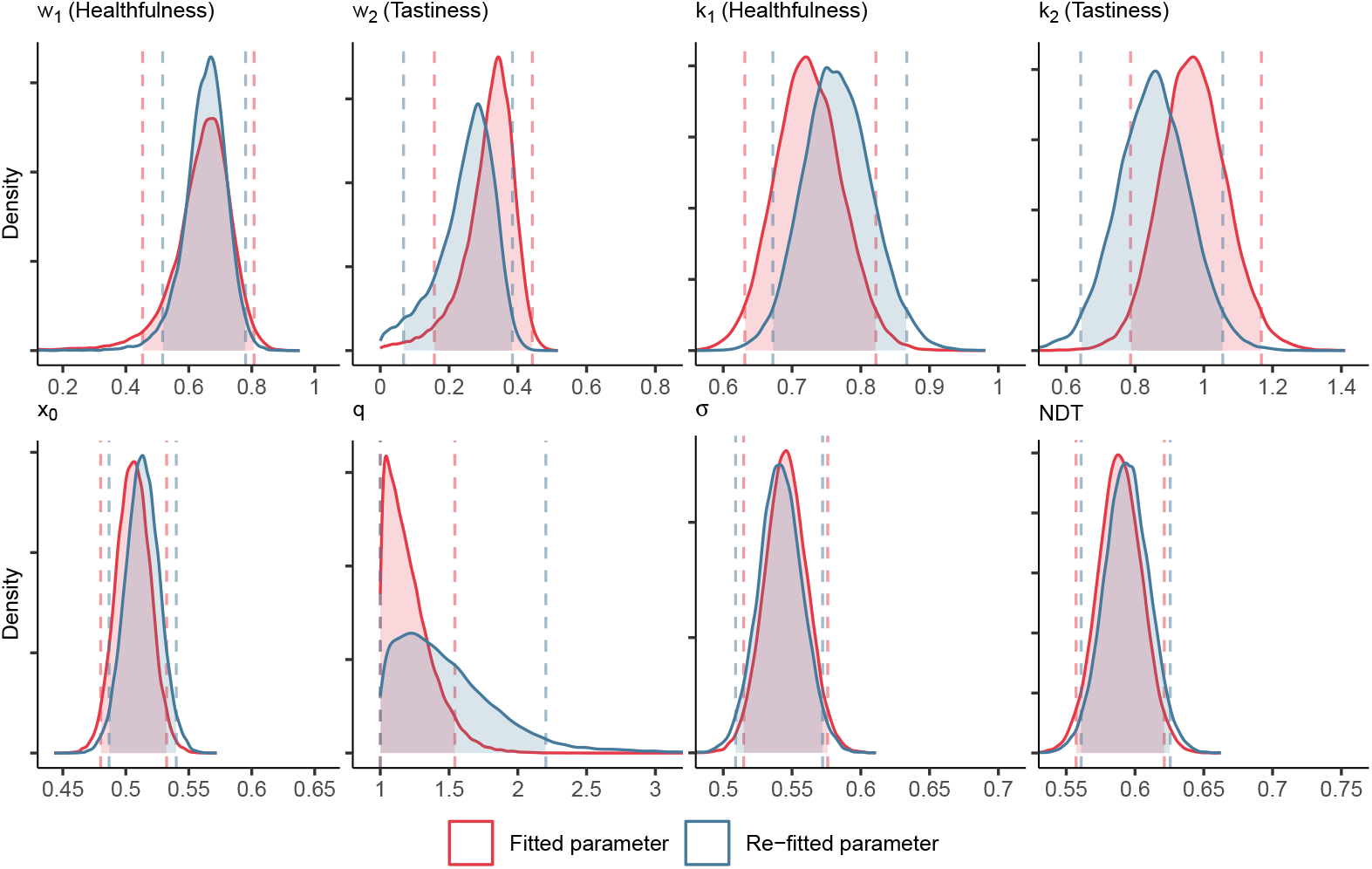
Parameter recovery for the nonlinear DDM with loss aversion in the Food2 dataset.

**Fig. S15.**
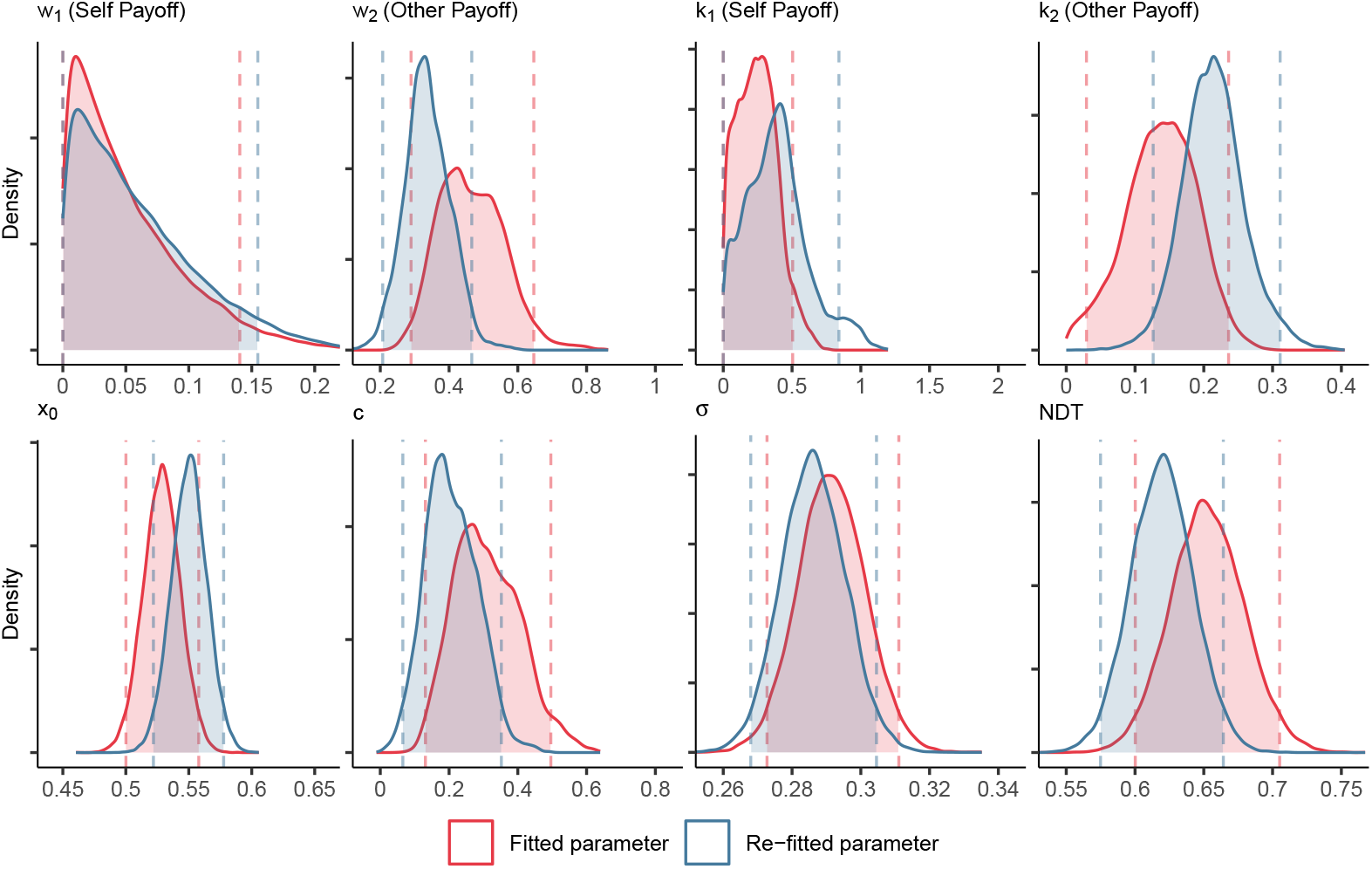
Parameter recovery for the nonlinear DDM in the Social1 dataset’s “Time-free” condition.

**Fig. S16.**
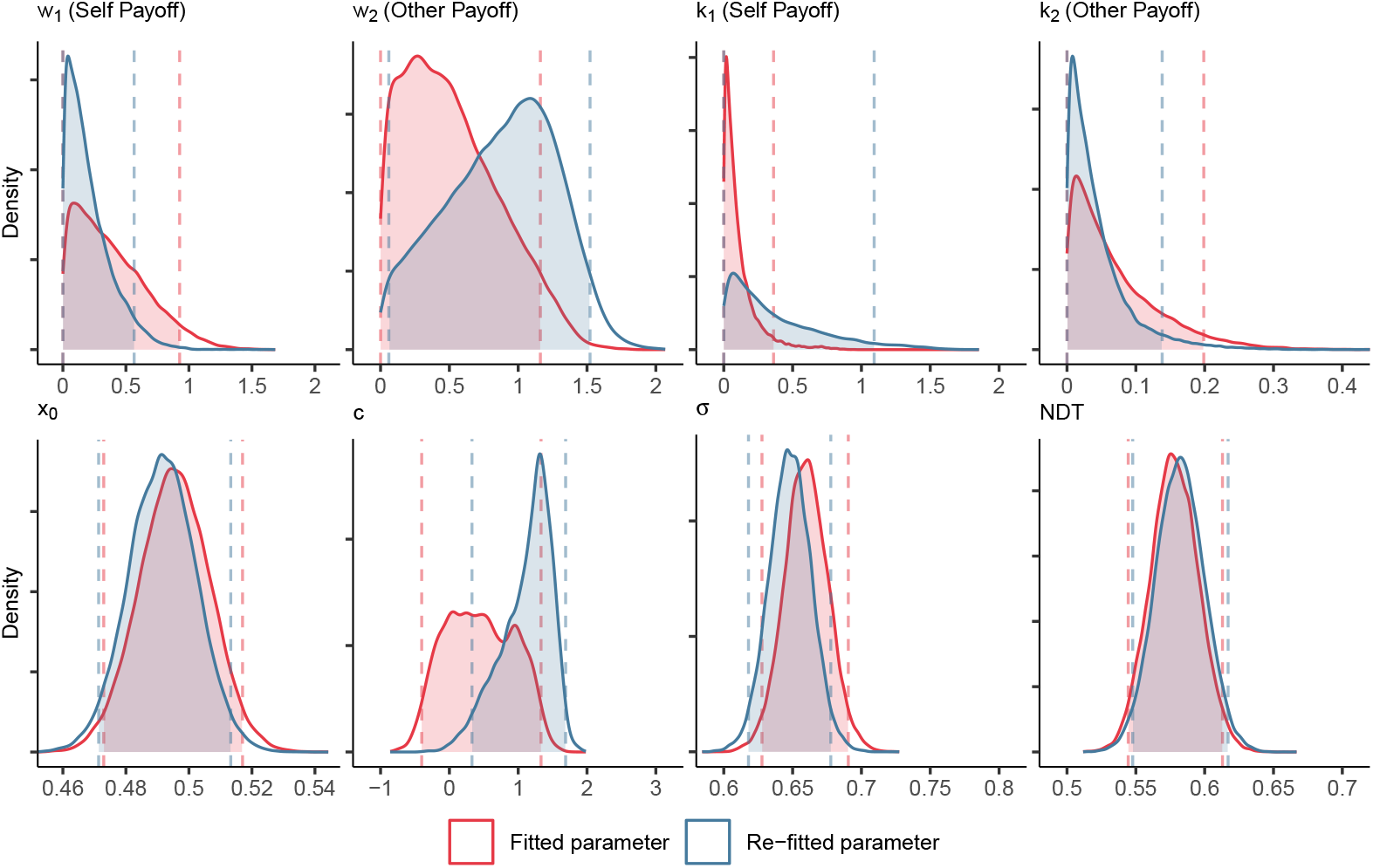
Parameter recovery for the nonlinear DDM in the Social1 dataset’s “Time-pressure” condition.

**Fig. S17.**
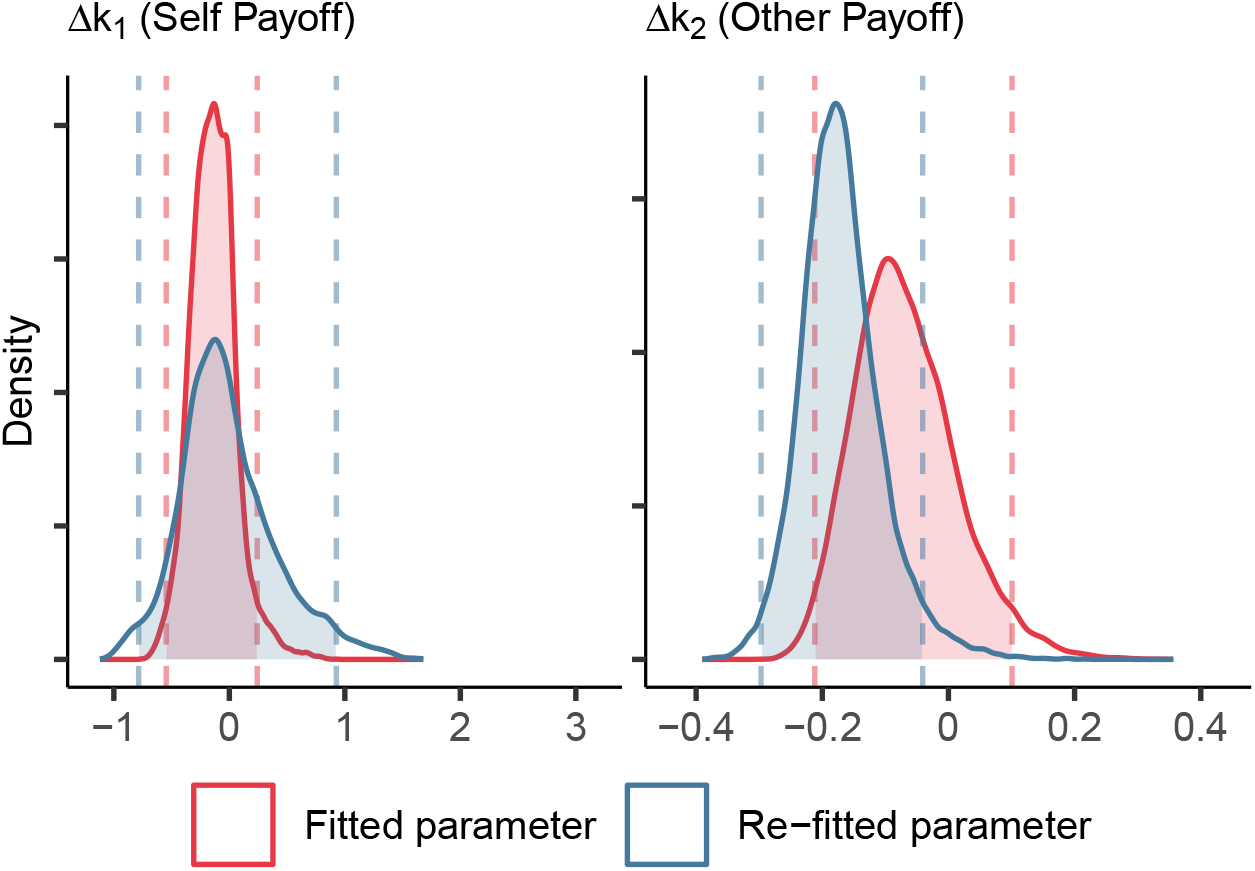
Parameter recovery for *k*_1_ and *k*_2_ parameters’s difference between “Time-pressure” condition and “Time-free” condition of the Social1 dataset.

**Fig. S18.**
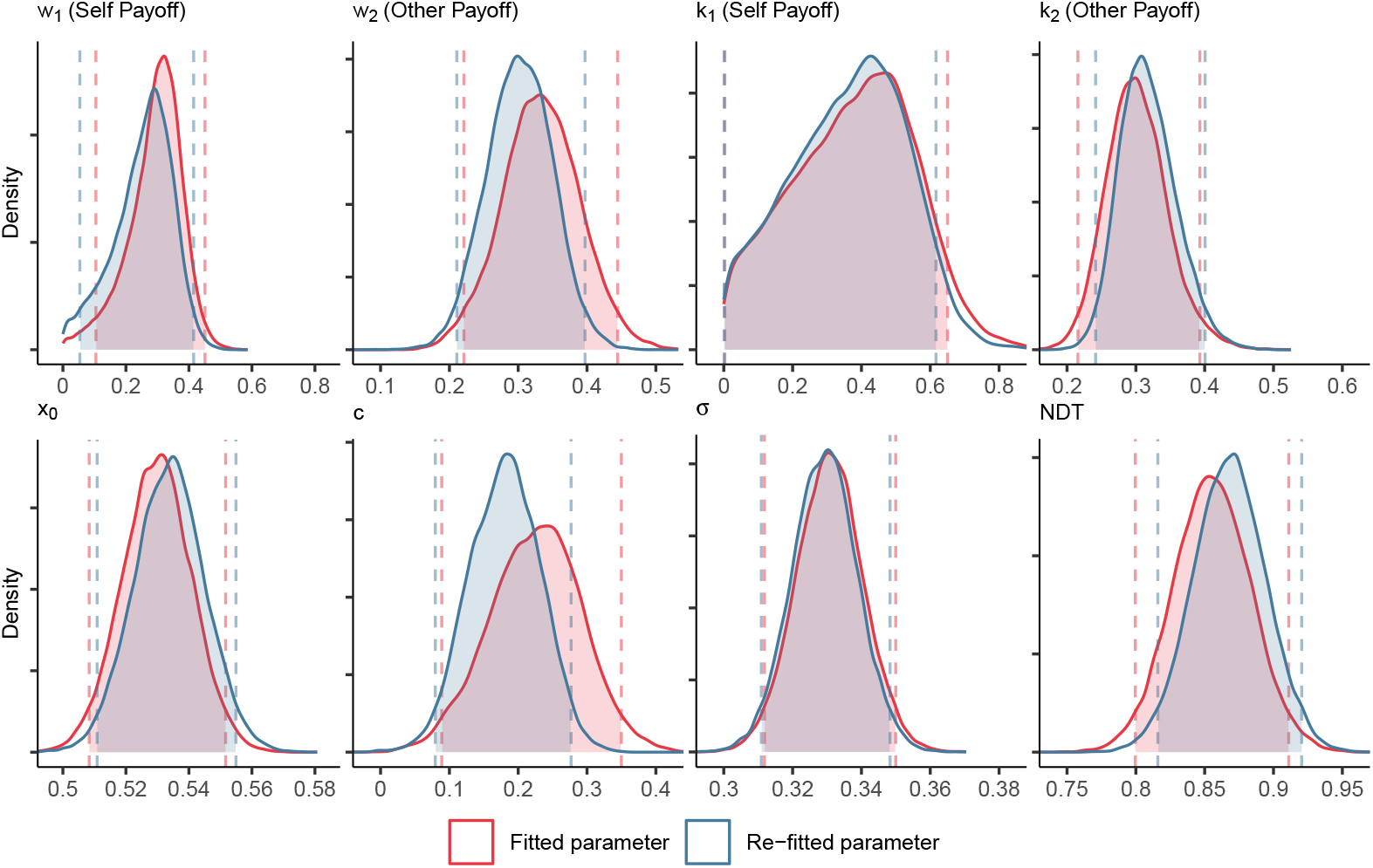
Parameter recovery for the nonlinear DDM in the Social2 dataset’s “Time-free” condition.

**Fig. S19.**
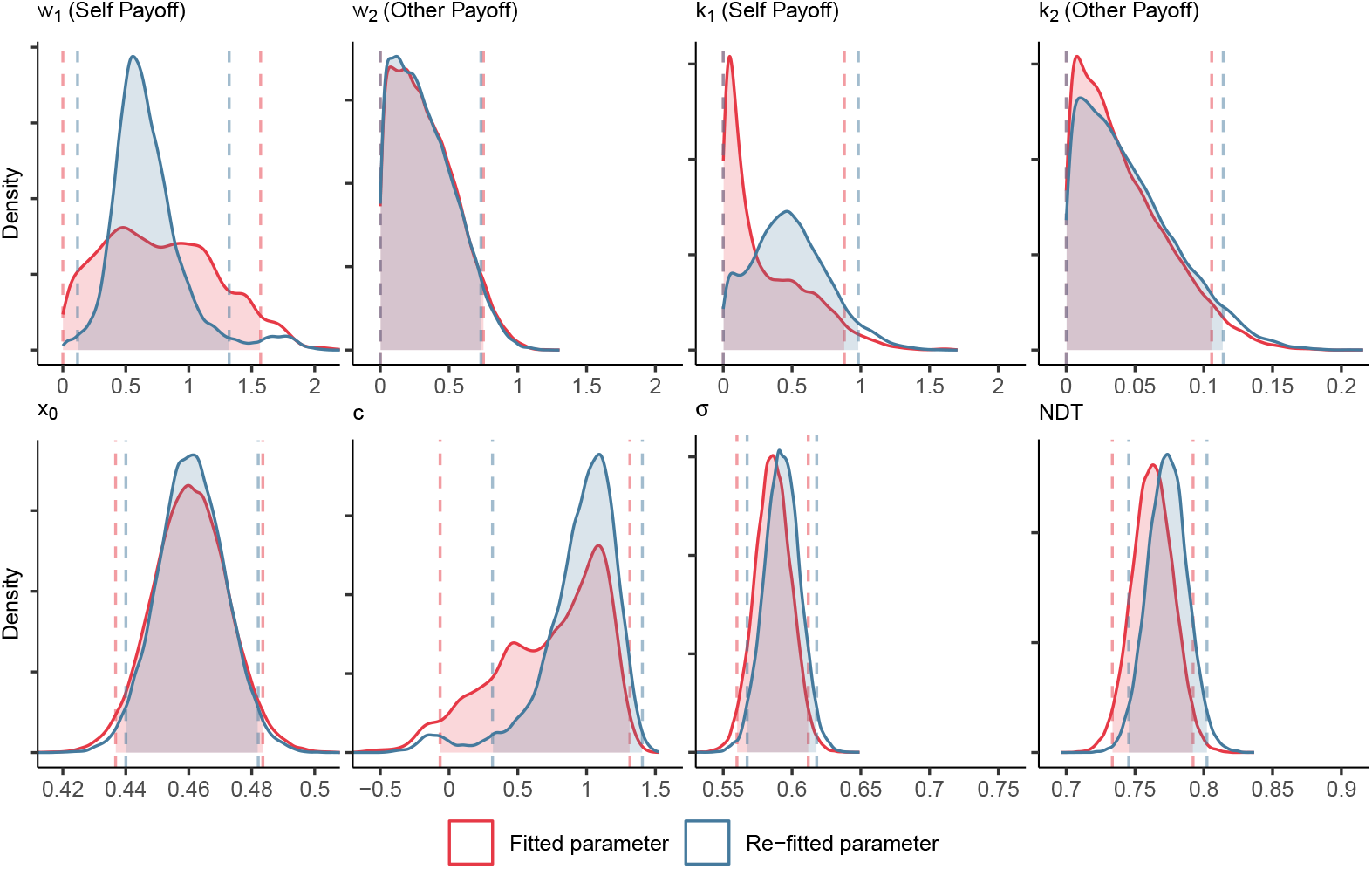
Parameter recovery for the nonlinear DDM in the Social2 dataset’s “Time-pressure” condition.

**Fig. S20.**
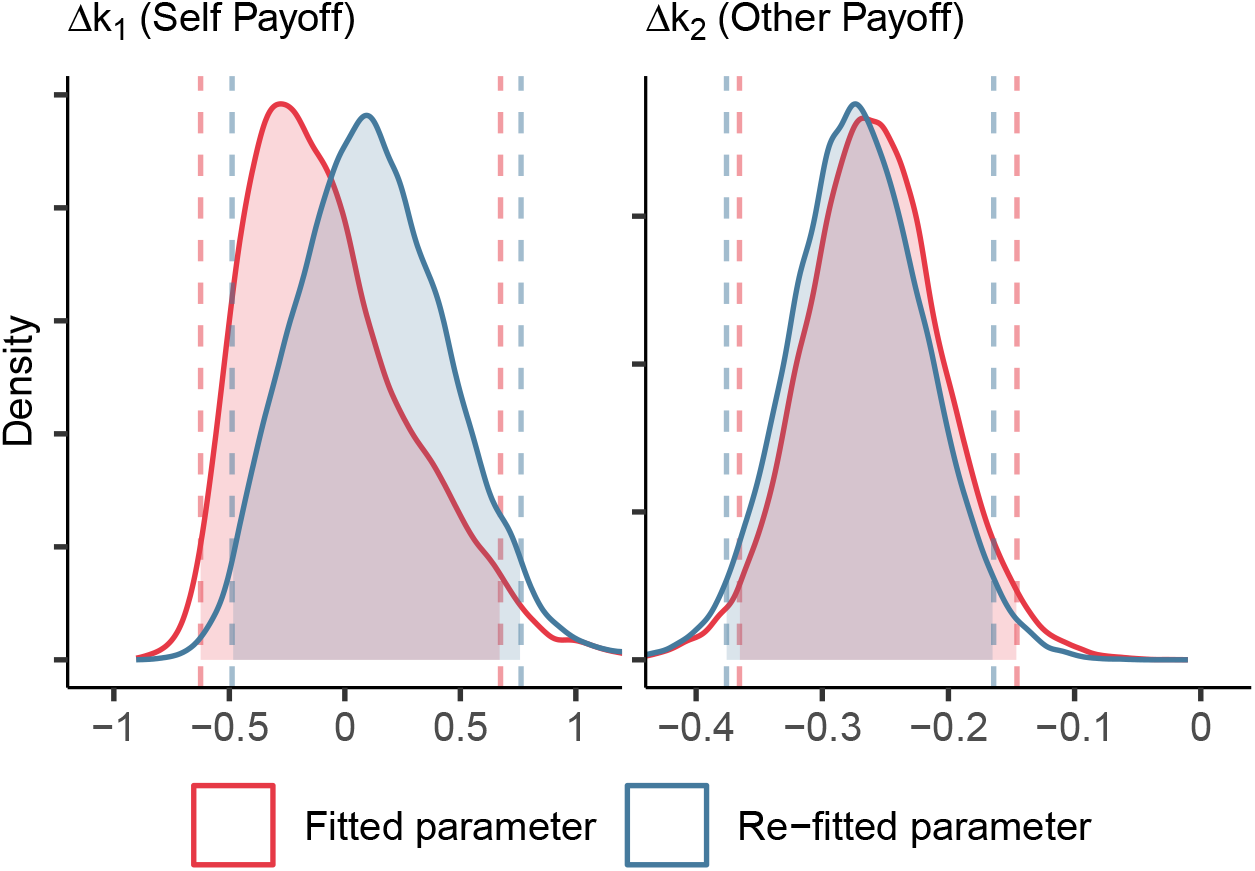
Parameter recovery for *k*_1_ and *k*_2_ parameters’s difference between “Time-pressure” condition and “Time-free” condition of the Social2 dataset.

### Comparison of homoscedastic and heteroscedastic model fits

The following table shows model comparison results for homoscedastic (*m*_*homo*_) and heteroscedastic (*m*_*hetero*_) models for modeling attribute value difference estimations based on AIC, BIC, and likelihood ratio test.

**Table S8.**
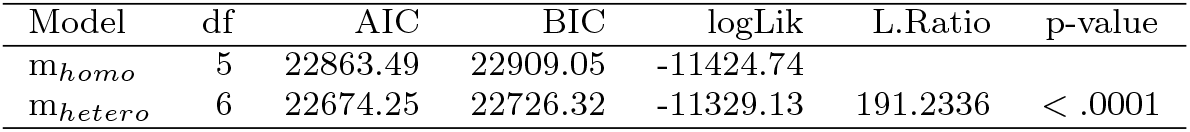
Model comparison (homoscedastic vs. heteroscedastic) for attribute value difference estimation.

| Model | df | AIC | BIC | logLik | L.Ratio | p-value |
| --- | --- | --- | --- | --- | --- | --- |
| $m_{homo}$ | 5 | 22863.49 | 22909.05 | -11424.74 | | |
| $m_{hetero}$ | 6 | 22674.25 | 22726.32 | -11329.13 | 191.2336 | < .0001 |

### Empirical evidence supporting the negligible bias–bias term

**Table S9.**
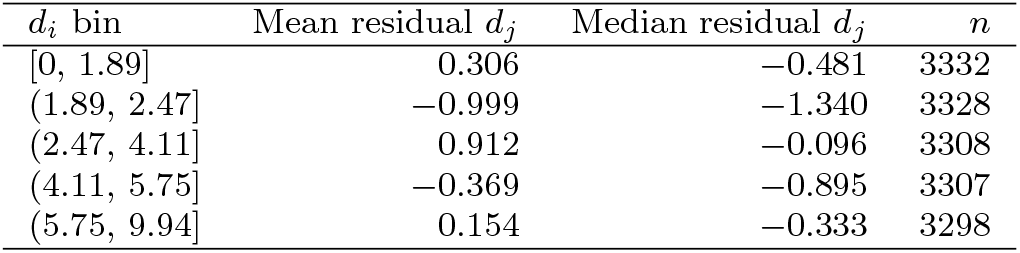
In Food1 dataset, residual statistics for *d*_*j*_ (healthfulness difference between tastier option and less tasty option) around the fitted linear trade-off line *d*_*j*_ = *α* + *βd*_*i*_, where *d*_*i*_ is tastiness difference. Residuals are small, supporting the approximation *χ*_*i*_(*d*_*i*_) ≈ 0.

**Table S10.**
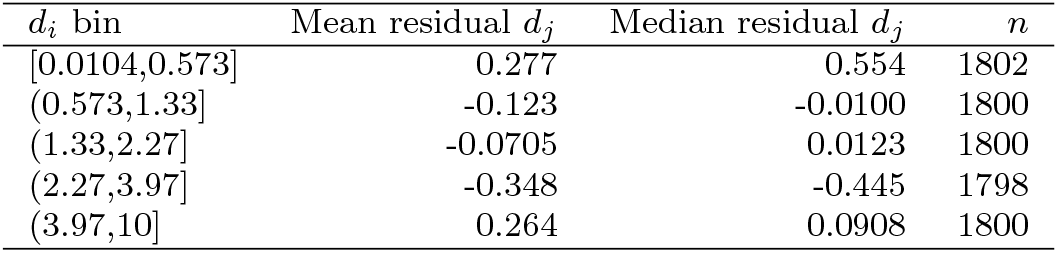
In Food2 dataset, residual statistics for *d*_*j*_ (healthfulness difference between tastier option and less tasty option) around the fitted linear trade-off line *d*_*j*_ = *α* + *βd*_*i*_, where *d*_*i*_ is tastiness difference. Residuals are small, supporting the approximation *χ*_*i*_(*d*_*i*_) ≈ 0.

**Table S11.**
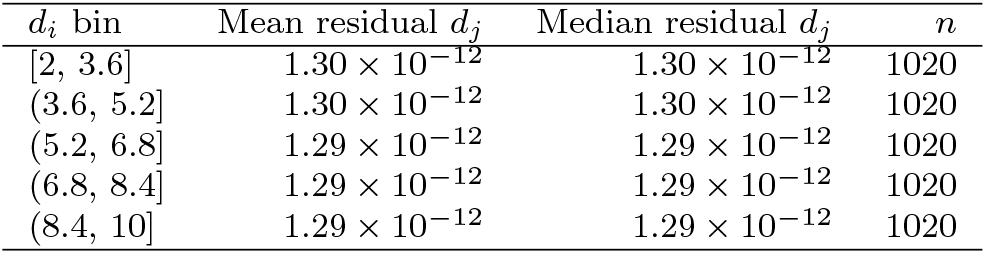
In Social1 dataset, residual statistics for *d*_*j*_ (own’s payoff difference between less prosocial option and more prosocial option) around the fitted linear trade-off line *d*_*j*_ = *α* + *βd*_*i*_, where *d*_*i*_ is other’s payoff difference. Residuals are near zero and symmetric, supporting the approximation *χ*_*i*_(*d*_*i*_) ≈ 0.

**Fig. S21.**
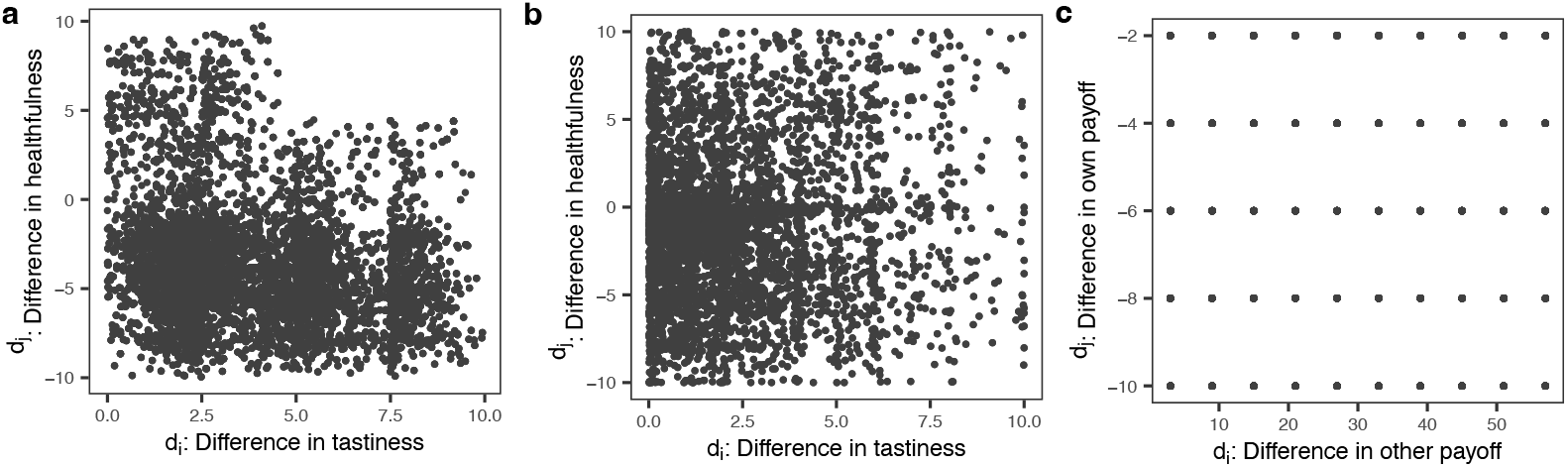
Symmetry of paired attribute differences supports the negligible cross-term assumption. Joint distributions of paired attribute differences (*d*_*j*_ versus *d*_*i*_) used to assess whether the cross-term *χ*_*i*_(*d*_*i*_) is small in the two-attribute case. (a-b) Food-choice datasets (Food1 and Food2): *d*_*i*_ denotes the difference in tastiness and *d*_*j*_ the difference in healthfulness, each computed as the tastier option minus the less tasty option. (c) Social-choice datasets: *d*_*i*_ denotes the difference in other payoff and *d*_*j*_ the difference in self payoff, each computed as the prosocial option minus the selfish option. Across datasets, conditional distributions of *d*_*j*_ given *d*_*i*_ are approximately centered and symmetric, supporting the approximation that *χ*_*i*_(*d*_*i*_) is negligible under the model’s symmetry assumptions.

## References

[1] Polanía, R., Woodford, M., Ruff, C.C.: Efficient coding of subjective value. Nature Neuroscience 22(1), 134–142 (2019)

[2] Keeney, R.L., Raiffa, H.: Decisions with Multiple Objectives: Preferences and Value Trade-offs. Cambridge university press, ??? (1993)

[3] Payne, J.W., Bettman, J.R., Johnson, E.J.: Adaptive strategy selection in decision making. 14(3), 534–552. Publisher: American Psychological Association

[4] Hutcherson, C.A., Bushong, B., Rangel, A.: A Neurocomputational Model of Altruistic Choice and Its Implications. Neuron 87(2), 451–462 (2015)

[5] Chen, F., Krajbich, I.: Biased sequential sampling underlies the effects of time pressure and delay in social decision making. Nature Communications 9(1), 3557 (2018)

[6] Maier, S.U., Raja Beharelle, A., Polanía, R., Ruff, C.C., Hare, T.A.: Dissociable mechanisms govern when and how strongly reward attributes affect decisions. Nature Human Behaviour 4(9), 949–963 (2020)

[7] Bhatia, S., Loomes, G., Read, D.: Establishing the laws of preferential choice behavior. Judgment and Decision Making 16(6), 1324–1369 (2021)

[8] Usher, M., McClelland, J.L.: Loss Aversion and Inhibition in Dynamical Models of Multialternative Choice. Psychological Review 111(3), 757–769 (2004)

[9] Chorus, C.G., Arentze, T.A., Timmermans, H.J.P.: A Random Regret-Minimization model of travel choice. Transportation Research Part B: Method-ological 42(1), 1–18 (2008)

[10] Scheibehenne, B., Miesler, L., Todd, P.M.: Fast and frugal food choices: Uncovering individual decision heuristics. Appetite 49(3), 578–589 (2007)

[11] Roering, K.J., Boush, D.M., Shipp, S.H., et al.: Factors that shape eating patterns: A consumer behavior perspective. What is America eating, 72–84 (1986)

[12] Sims, C.R.: Rate–distortion theory and human perception. Cognition 152, 181–198 (2016)

[13] Cheyette, S.J., Piantadosi, S.T.: A unified account of numerosity perception. Nature human behaviour 4(12), 1265–1272 (2020)

[14] Ortega, P.A., Braun, D.A.: Thermodynamics as a theory of decision-making with information-processing costs. Proceedings of the Royal Society A: Mathematical, Physical and Engineering Sciences 469(2153), 20120683 (2013)

[15] Lieder, F., Griffiths, T.L.: Resource-rational analysis: Understanding human cognition as the optimal use of limited computational resources. Behavioral and brain sciences 43, 1 (2020)

[16] Schaffner, J., Bao, S.D., Tobler, P.N., Hare, T.A., Polania, R.: Sensory perception relies on fitness-maximizing codes. Nature Human Behaviour 7(7), 1135–1151 (2023)

[17] Zhang, H., Ren, X., Maloney, L.T.: The bounded rationality of probability distortion. Proceedings of the National Academy of Sciences 117(36), 22024–22034 (2020)

[18] Griffiths, T.L., Lieder, F., Goodman, N.D.: Rational use of cognitive resources: Levels of analysis between the computational and the algorithmic. Topics in cognitive science 7(2), 217–229 (2015)

[19] Cheyette, S.J., Wu, S., Piantadosi, S.T.: Limited information-processing capacity in vision explains number psychophysics. Psychological Review 131(4), 891–904 (2024)

[20] Laughlin, S.B.: Energy as a constraint on the coding and processing of sensory information. Current opinion in neurobiology 11(4), 475–480 (2001)

[21] Polanía, R., Burdakov, D., Hare, T.A.: Rationality, preferences, and emotions with biological constraints: it all starts from our senses. Trends in Cognitive Sciences 28(3), 264–277 (2024)

[22] Cover, T.M.: Elements of Information Theory. John Wiley & Sons, ??? (1999)

[23] Dehaene, S.: Cross-linguistic regularities in the frequency of number words. Cognition 43(1), 1–29 (1992)

[24] Piantadosi, S.T.: A rational analysis of the approximate number system. Psychonomic Bulletin & Review 23(3), 877–886 (2016)

[25] Piantadosi, S.T., Cantlon, J.F.: True Numerical Cognition in the Wild. Psychological Science 28(4), 462–469 (2017)

[26] Clithero, J.A.: Improving out-of-sample predictions using response times and a model of the decision process 148, 344–375

[27] Pedersen, M.L., Frank, M.J., Biele, G.: The drift diffusion model as the choice rule in reinforcement learning 24(4), 1234–1251

[28] Vehtari, A., Gelman, A., Gabry, J.: Practical Bayesian model evaluation using leave-one-out cross-validation and WAIC. Statistics and Computing 27(5), 1413–1432 (2017)

[29] Engel, C.: Dictator games: a meta study. Experimental Economics 14(4), 583–610 (2011)

[30] Lindig-León, C., Kaur, N., Braun, D.A.: From bayes-optimal to heuristic decision-making in a two-alternative forced choice task with an information-theoretic bounded rationality model. Frontiers in Neuroscience 16, 906198 (2022)

[31] Ortega, P.A., Stocker, A.A.: Human decision-making under limited time. Advances in Neural Information Processing Systems 29 (2016)

[32] Wu, C.M., Schulz, E., Pleskac, T.J., Speekenbrink, M.: Time pressure changes how people explore and respond to uncertainty. Scientific reports 12(1), 4122 (2022)

[33] Attneave, F.: Some informational aspects of visual perception. Psychological review 61(3), 183 (1954)

[34] Barlow, H.B., et al.: Possible principles underlying the transformation of sensory messages. Sensory communication 1(01), 217–233 (1961)

[35] Berger, T.: Rate-distortion theory. Wiley Encyclopedia of Telecommunications (2003)

[36] Wei, X.-X., Stocker, A.A.: A Bayesian observer model constrained by efficient coding can explain ‘anti-Bayesian’ percepts. Nature Neuroscience 18(10), 1509–1517 (2015)

[37] Wei, X.-X., Stocker, A.A.: Lawful relation between perceptual bias and discriminability. Proceedings of the National Academy of Sciences 114(38), 10244–10249 (2017)

[38] Hahn, M., Wei, X.-X.: A unifying theory explains seemingly contradictory biases in perceptual estimation. Nature Neuroscience 27(4), 793–804 (2024)

[39] Heng, J.A., Woodford, M., Polania, R.: Efficient sampling and noisy decisions. Elife 9, 54962 (2020)

[40] Heng, J.A., Woodford, M., Polania, R.: Efficient numerosity estimation under limited time. PLOS Computational Biology 21(3), 1012790 (2025)

[41] Prat-Carrabin, A., Woodford, M.: Efficient coding of numbers explains decision bias and noise. Nature Human Behaviour 6(8), 1142–1152 (2022)

[42] Cheyette, S.J., Wu, S., Piantadosi, S.T.: Limited information-processing capacity in vision explains number psychophysics. Psychological Review 131(4), 891 (2024)

[43] Sims, C.R., Jacobs, R.A., Knill, D.C.: An ideal observer analysis of visual working memory. Psychological review 119(4), 807 (2012)

[44] Sims, C.R.: The cost of misremembering: Inferring the loss function in visual working memory. Journal of vision 15(3), 2–2 (2015)

[45] Sims, C.R.: Efficient coding explains the universal law of generalization in human perception. Science 360(6389), 652–656 (2018)

[46] Khaw, M.W., Li, Z., Woodford, M.: Cognitive imprecision and small-stakes risk aversion. The review of economic studies 88(4), 1979–2013 (2021)

[47] Frydman, C., Jin, L.J.: Efficient coding and risky choice. The Quarterly Journal of Economics 137(1), 161–213 (2022)

[48] Bedi, S., Hollander, G., Ruff, C.C.: Probability weighting arises from boundary repulsions of cognitive noise. bioRxiv, 2025–09 (2025)

[49] Woodford, M.: Prospect theory as efficient perceptual distortion. American Economic Review 102(3), 41–46 (2012)

[50] Barretto-García, M., De Hollander, G., Grueschow, M., Polanía, R., Wood-ford, M., Ruff, C.C.: Individual risk attitudes arise from noise in neurocognitive magnitude representations. Nature Human Behaviour 7(9), 1551–1567 (2023)

[51] Hsu, M., Krajbich, I., Zhao, C., Camerer, C.F.: Neural Response to Reward Anticipation under Risk Is Nonlinear in Probabilities. The Journal of Neuroscience 29(7), 2231–2237 (2009)

[52] Zalocusky, K.A., Ramakrishnan, C., Lerner, T.N., Davidson, T.J., Knutson, B., Deisseroth, K.: Nucleus accumbens D2R cells signal prior outcomes and control risky decision-making. Nature 531(7596), 642–646 (2016)

[53] Schaffner, J., Bao, S.D., Tobler, P.N., Hare, T.A., Polania, R.: Sensory perception relies on fitness-maximizing codes. Nature Human Behaviour 7(7), 1135–1151 (2023)

[54] Leong, Y.C., Hughes, B.L., Wang, Y., Zaki, J.: Neurocomputational mechanisms underlying motivated seeing. Nature human behaviour 3(9), 962–973 (2019)

[55] Banerjee, A., Parente, G., Teutsch, J., Lewis, C., Voigt, F.F., Helmchen, F.: Value-guided remapping of sensory cortex by lateral orbitofrontal cortex. Nature 585(7824), 245–250 (2020)

[56] Liu, Y., Xin, Y., Xu, N.-l.: A cortical circuit mechanism for structural knowledge-based flexible sensorimotor decision-making. Neuron 109(12), 2009–2024 (2021)

[57] Amasino, D.R., Sullivan, N.J., Kranton, R.E., Huettel, S.A.: Amount and time exert independent influences on intertemporal choice. Nature Human Behaviour 3(4), 383–392 (2019)

[58] Fellows, L.K.: Deciding how to decide: ventromedial frontal lobe damage affects information acquisition in multi-attribute decision making. Brain 129(4), 944–952 (2006)

[59] Yang, X., Krajbich, I.: A dynamic computational model of gaze and choice in multi-attribute decisions. Psychological Review (2022)

[60] Perkins, A.Q., Gillis, Z.S., Rich, E.L.: Multiattribute Decision-making in Macaques Relies on Direct Attribute Comparisons. Journal of Cognitive Neuro-science 36(9), 1879–1897 (2024)

[61] Perkins, A.Q., Rich, E.L.: Orbitofrontal cortex computes gaze-dependent comparisons between attributes rather than integrated values. PLOS Biology 23(8), 3003281 (2025)

[62] Setogawa, T., Mizuhiki, T., Matsumoto, N., Akizawa, F., Kuboki, R., Rich-mond, B.J., Shidara, M.: Neurons in the monkey orbitofrontal cortex mediate reward value computation and decision-making. Communications Biology 2(1), 126 (2019)

[63] Magrabi, A., Ludwig, V.U., Stoppel, C.M., Paschke, L.M., Wisniewski, D., Heek-eren, H.R., Walter, H.: Dynamic computation of value signals via a common neural network in multi-attribute decision-making. Social Cognitive and Affective Neuroscience 17(7), 683–693 (2022)

[64] González-Vallejo, C.: Making trade-offs: a probabilistic and context-sensitive model of choice behavior. Psychological Review 109(1), 137 (2002)

[65] González-Vallejo, C., Reid, A.A., Schiltz, J.: Context effects: The proportional difference model and the reflection of preference. Journal of Experimental Psychology: Learning, Memory, and Cognition 29(5), 942 (2003)

[66] Scholten, M., Read, D.: The psychology of intertemporal tradeoffs. Psychological review 117(3), 925 (2010)

[67] Mezzadri, G., Woodford, M.: Sources of imprecision in integrated value comparisons. Available at SSRN 6017358 (2025)

[68] Trueblood, J.S., Brown, S.D., Heathcote, A.: The multiattribute linear ballistic accumulator model of context effects in multialternative choice. Psychological Review 121(2), 179–205 (2014)

[69] Bordalo, P., Gennaioli, N., Shleifer, A.: Salience and Consumer Choice. Journal of Political Economy 121(5), 803–843 (2013)

[70] Landry, P., Webb, R.: Pairwise normalization: A neuroeconomic theory of multi-attribute choice. Journal of Economic Theory 193, 105221 (2021)

[71] Soltani, A., De Martino, B., Camerer, C.: A Range-Normalization Model of Context-Dependent Choice: A New Model and Evidence. PLoS Computational Biology 8(7), 1002607 (2012)

[72] Hunt, L.T., Dolan, R.J., Behrens, T.E.J.: Hierarchical competitions subserving multi-attribute choice. Nature Neuroscience 17(11), 1613–1622 (2014)

[73] Roe, R.M., Busemeyer, J.R., Townsend, J.T.: Multialternative decision field theory: A dynamic connectionst model of decision making. Psychological Review 108(2), 370–392 (2001)

[74] Rramani, Q., Krajbich, I., Enax, L., Brustkern, L., Weber, B.: Salient nutrition labels shift peoples’ attention to healthy foods and exert more influence on their choices. Nutrition Research 80, 106–116 (2020)

[75] Boyd, S.P., Vandenberghe, L.: Convex Optimization. Cambridge university press, ??? (2004)

[76] Nesterov, Y.: Introductory Lectures on Convex Optimization: A Basic Course vol. 87. Springer, ??? (2013)

[77] Cheyette, S.J., Piantadosi, S.T.: A unified account of numerosity perception. Nature Human Behaviour 4(12), 1265–1272 (2020)

[78] Ska-lbania, J., Tanajewski, L., Furtak, M., Hare, T.A., Wypych, M.: Pre-choice midbrain fluctuations affect self-control in food choice: A functional magnetic resonance imaging (fMRI) study. Cognitive, Affective, & Behavioral Neuroscience (2024)

[79] Chen, F., Zhu, Z., Shen, Q., Krajbich, I., Hare, T.A.: Intrachoice Dynamics Shape Social Decisions. Management Science 70(2), 1137–1153 (2024)

[80] Plummer, M.: JAGS: A program for analysis of Bayesian graphical models using Gibbs sampling

[81] Sokol-Hessner, P., Hsu, M., Curley, N.G., Delgado, M.R., Camerer, C.F., Phelps, E.A.: Thinking like a trader selectively reduces individuals’ loss aversion. Proceedings of the National Academy of Sciences 106(13), 5035–5040 (2009)

